# Less, but more: new insights from appendicularians on chordate *Fgf* evolution and the divergence of tunicate lifestyles

**DOI:** 10.1101/2024.08.30.610304

**Authors:** Gaspar Sánchez-Serna, Jordi Badia-Ramentol, Paula Bujosa, Alfonso Ferrández-Roldán, Nuria P. Torres-Águila, Marc Fabregà-Torrus, Johannes N. Wibisana, Michael J. Mansfield, Charles Plessy, Nicholas M. Luscombe, Ricard Albalat, Cristian Cañestro

## Abstract

The impact of gene loss on the divergence of taxa and the generation of evolutionary innovations is a fundamental aspect of Evolutionary Biology that remains unclear. Here, using the evolution of the Fibroblast Growth Factors (FGFs) in appendicularians as a case study, we investigate how gene losses have influenced the evolution of chordates, especially the divergence among tunicates. Our work reveals an unprecedented case of massive losses of all *Fgf* gene subfamilies, except for the *Fgf9/16/20* and *Fgf11/12/13/14*, which in turn suffered two bursts of gene duplications. Phylogenetic inferences and genomic analyses of gene synteny conservation, gene architecture, alternative splicing and protein 3D-structure have allowed us to reconstruct the history of appendicularian *Fgf* genes in the context of chordate evolution, providing compelling evidence supporting the paracrine secreting functions and the intracellular functions of the *Fgf9/16/20* and *Fgf11/12/13/14* subfamilies, respectively. Exhaustive analysis of developmental *Fgf* expression in *Oikopleura dioica* as a model for appendicularians reveals a paradigmatic case of what could be referred as “less, but more”, providing a conceptual evolutionary framework characterized by four associated evolutionary patterns: conservation of ancestral *Fgf* expression domains; function shuffling between paralogs upon gene loss; innovation of new expression domains after the bursts of *Fgf* duplications; and the extinction of *Fgf* functions linked to gene losses. The findings of this work allow us to formulate novel hypotheses about the potential impact of losses and duplications of *Fgf* genes on the transition from an ancestral ascidian-like biphasic lifestyle to a fully free-living style of appendicularians. These hypotheses include the massive co-option of *Fgf* genes for the patterning of the oikoblast responsible of the house architecture, and for the development of the tail fin; the recruitment of *Fgf11/12/13/14* genes into the evolution of a new mouth, and their role modulating neuronal excitability; the evolutionary innovation of an “anterior tail” FGF signaling mesodermal source upon the loss of retinoic acid signaling; and the potential link between the loss of *Fgf7/10/22* and *Fgf8/17/18* and the loss of drastic metamorphosis, mesenchymal cells and lack of tail absorption in appendicularians, in contrast to ascidians.

## Introduction

The bloom of sequenced genomes has revealed that gene losses are pervasive and prevalent over gene gains throughout the tree of life, leaving little doubt of their great potential as a key evolutionary force that can generate adaptive phenotypic diversity (Krylov et al. 2003; Albalat and Cañestro 2016; Fernández and Gabaldón 2020; Guijarro-Clarke et al. 2020; Helsen et al. 2020; Xu and Guo 2020). There are many paradigmatic examples of gene losses that have been key for the evolution of adaptations in certain species, under what is known as the “less is more” hypothesis (Olson 1999). Examples of adaptive gene losses include positively selected null-mutations in certain receptors that provide resistance to malaria and HIV in humans (Novembre et al. 2005; Hodgson et al. 2014), the loss of gluconeogenic muscle enzyme that allowed the evolution of true hovering flight in hummingbirds (Osipova et al. 2023), and many gene losses that facilitated the reconquest of aquatic and air environments in mammals (Sharma et al. 2018). However, how evolutionary processes can drive the loss of essential genes without carrying an important detrimental load remains enigmatic. The loss of a gene often does not come as an isolated event, but it is accompanied by the co-elimination of other genes that are functionally linked to a distinctive pathway (Albalat and Cañestro 2016). Moreover, within a given gene family, the loss of some members is often accompanied by the duplication of others, what increases the robustness of the genetic system and may lead to processes of function shuffling among paralogs that facilitate the events of gene loss (McClintock et al. 2001; Cañestro et al. 2009). To understand the impact of the loss of essential genes, such as those governing embryo development, it is necessary to study cases in which events of gene co-elimination and duplication can be related to the loss or survival of ancestral traits still present in sister groups, or even to the origin of evolutionary adaptations.

During recent years, the appendicularian tunicate *Oikopleura dioica* has become an attractive animal model to study the impact of gene loss on the evolution of developmental mechanisms in our own phylum, the chordates (Ferrández-Roldán et al. 2019). For instance, the discovery of the deconstruction of the cardiopharyngeal gene regulatory network in *O. dioica* has been key to understanding the adaptive evolution of the heart, facilitating the transition from an ancestral biphasic ascidian-like lifestyle to the complete free-living lifestyle in appendicularians (Ferrández-Roldán et al. 2021). *O. dioica* also appears to have suffered a large number of gene losses affecting important signaling pathways, such as the wingless (Wnt) and retinoic acid (RA) signaling pathways, that play fundamental roles in axial patterning and cell differentiation during embryo development in chordates (Martí-Solans et al. 2016; Martí-Solans et al. 2021). *O. dioica* has the minimal Wnt repertoire found among chordates, where only four out of the thirteen Wnt families are present. *O. dioica* also stands as the only chordate known to date that can be considered an evolutionary knockout model for RA signaling. The antagonistic action of the RA and the fibroblast growth factor (FGF) signaling pathways is well known in vertebrates during axial patterning, where it regulates for instance the expression of *Hox* genes (Diez del Corral and Storey 2004; Wilson et al. 2009). The fact that this antagonistic action has also been described in ascidians led to conclude that the RA-FGF antagonism was already present in the last common ancestor of olfactores (this is vertebrates + tunicates) (Pasini et al. 2012). Moreover, its absence in cephalochordates led to suggest that the evolutionary innovation of this RA-FGF antagonism could have facilitated the evolutionary origin and radiation of olfactores (Bertrand et al. 2015). The loss of RA signaling and the drastic reduction of Wnt families in *O. dioica* makes the evolution of FGF signaling in this species particularly intriguing. Its study can contribute to a better understanding of how living beings manage the loss of important genes while maintaining similar morphologies (i.e., the inverse paradox in Evo-Devo) (Cañestro et al. 2007), and to correlating gene loss events with evolutionary adaptations of the lifestyle of certain groups of organisms.

Fibroblast growth factors form a family of signaling proteins that emerged concomitantly with the origin of Eumetazoans and have been vastly conserved during animal evolution, regulating a plethora of important biological processes such as cell proliferation, migration, or differentiation during embryonic development and adult tissue homeostasis (Bertrand et al. 2014; Teven et al. 2014). In general, Fgfs are small proteins characterized by a conserved FGF core homology domain of 120-130 amino acids disposed in a β-trefoil topology (Plotnikov et al. 2001). The FGF domain contains essential motifs for binding both heparin and the extracellular region of the Fgf receptors on the cell surface, forming a dimeric ternary complex that can trigger canonical signal transduction cascades inside the cells (Schlessinger et al. 2000). Outside the FGF domain, sequences are not conserved among different subfamilies, and some *Fgf* genes have independently evolved extended N- or C-terminal regions that are not homologous (Popovici et al. 2005). Consequently, protein alignments between distant Fgf subfamilies are not possible outside the FGF domain. Some of these extended regions of variable lengths often include signal peptides (SP) and nuclear localization signals (NLS) (Coulier et al. 1997). The SP is a short hydrophobic region that includes the first ∼15-30 residues of a protein at its N-terminus and is important for the secretion of proteins outside the cell (Owji et al. 2018). In principle, Fgfs that lack an SP remain intracellular, although alternative secretion mechanisms that enable these Fgfs to perform paracrine signaling functions have also been described (Revest et al. 2000; Miyakawa and Imamura 2003; Schäfer et al. 2004; Kirov et al. 2012). Moreover, there is an increasing body of evidence for intracellular functions and interacting partners for non-secreted Fgfs (Goldfarb 2001; Schoorlemmer and Goldfarb 2002; Olsnes et al. 2003; Goldfarb 2005; Wu et al. 2012; Sluzalska et al. 2021). In addition, the presence of NLS allows some Fgfs to migrate into the nucleus and interact with other transcription factors to modulate the expression of target genes (Antoine et al. 2005; Bryant and Stow 2005; Sheng et al. 2005; Popovici et al. 2006).

The high sequence variability among *Fgfs* and the short length of their conserved core have hindered the phylogenetic classification of this family (Popovici et al. 2005). Recent evolutionary reconstructions suggested the existence of eight *Fgf* subfamilies in chordates (namely *Fgf1/2*, *Fgf3*, *Fgf4/5/6*, *Fgf7/10/22*, *Fgf8/17/18/24*, *Fgf9/16/20*, *Fgf11/12/13/14* and *Fgf19/21/23*) (Oulion et al. 2012). It has been proposed that the basally divergent chordate amphioxus might possess the full catalogue of chordate *Fgfs*, with one single member for each of the eight subfamilies. This contrasts with the large number of *Fgfs* within each subfamily in vertebrates due to the various rounds of genome duplication that occurred during the evolution of different lineages (e.g. 19 in sarcopterygians, 22 in mammals, 23 in chicken and 27 in zebrafish) (Dehal and Boore 2005; Bertrand et al. 2011; Oulion et al. 2012). In ascidian tunicates, seven *Fgf* genes have been found representing at least six of the eight chordate subfamilies, and suggesting that *Fgf1/2* and *Fgf3* were lost during the evolution of the lineage leading to ascidians, at the same time that a novel *Fgf* member, *Fgf-L*, was innovated probably as a duplicate of *Fgf7/10/22* (Dehal and Boore 2005; Popovici et al. 2005; Oulion et al. 2012).

According to their functions, *Fgf* subfamilies have been classically classified into: (I) canonical Fgfs (i.e. *Fgf1/2*, *Fgf3*, *Fgf4/5/6*, *Fgf7/10/22*, *Fgf8/17/18/24*, *Fgf9/16/20*), which act as paracrine and autocrine signaling factors binding and activating the Fgf-receptor tyrosine kinases; (II) endocrine Fgfs (i.e. *Fgf19/21/23*), which bind the cofactor Klotho instead of heparin and act as long-distance signaling molecules in vertebrates, and (III) intracellular non-secreted Fgfs (i.e. *Fgf11/12/13/14*) with intracrine non-signaling functions that serve as cofactors to other proteins (reviewed in Ornitz and Itoh 2015; Ornitz and Itoh 2022).

In the present work, as a case study to better understand the impact of gene loss, we address the evolution of the FGF signaling pathway in appendicularian tunicates. The independent genome assemblies at the chromosome level of three different cryptic species of *O. dioica* from different parts of the globe (i.e. Barcelona, Osaka, and Okinawa) (Plessy et al. 2024) allow us to identify for the first time with high confidence the full catalogue of *Fgf* genes in an appendicularian species, and to describe its complete atlas of expression during embryonic and larval development. Our results reveal that only two out of the eight chordate *Fgf* subfamilies have survived in *O. dioica*, and that the presence of 10 *Fgfs* in this species is the result of several gene duplications and losses that occurred during the evolution of the appendicularian lineage. Our findings, moreover, allow us to discuss how the massive losses and burst of duplications affecting the *Fgf* family have impacted on the developmental mechanisms underlying the evolution of the appendicularian free-swimming lifestyle in this unique chordate, which functions as an evolutionary knockout for RA signaling. Finally, the results of this case study allow us to propose a “less, but more” scenario as an conceptual framework that facilitates a better understanding of the evolutionary impact of gene loss.

## Material and methods

### Laboratory culture of *Oikopleura dioica*

*O. dioica* specimens were acquired from animal colonies that have been maintained in our facility in the University of Barcelona for over five years. The founder individuals were originally obtained from the Mediterranean coast near Barcelona (Catalonia, Spain) and cultured as detailed in Martí-Solans et al. 2015 (Martí-Solans et al. 2015). This project did not raise any ethical concerns since the experimentation conducted on aquatic invertebrate animals does not fall under the regulations pertaining to animal experimentation, as stipulated in Real Decreto 223 14-3-1998 and Catalonia Ley 5/1995, DOGC2073,5172. Nonetheless, all experimental procedures adhered to the European Union (EU) guidelines for animal care and were formally approved by the Ethical Animal Experimentation Committee (CEEA-2009) of the University of Barcelona.

### Genome database searches, gene identification and phylogenetic analyses

*Fgf* genes in *O*. *dioica* were first identified in the reference database of *O. dioica* relative to the Norwegian population (http://oikoarrays.biology.uiowa.edu/Oiko/) (Danks et al. 2013) with BLASTp and tBLASTn, using as queries the Fgf protein sequences from the vertebrate *Homo sapiens* and the tunicate *Ciona robusta*, as well as several other chordates. The corresponding orthologs were then identified in the telomere-to-telomere genomic assemblies of the Barcelona and Okinawa *O. dioica* species, and in the superscaffolded version of the Osaka *O. dioica* cryptic species using tBLASTn (Plessy et al. 2024). In each cryptic species, the newly identified *Fgf* genes were used to search for further paralogs using tBLASTn to obtain the final *Fgf* catalogue. The final catalogue in Barcelona was confirmed searching for proteins containing the FGF domain (PF00167), retrieved from the Pfam database, with HMMscan against all the transcripts translated in all reading frames from a Barcelona *O. dioica* transcriptome assembly (Plessy et al. 2024).

*Fgf* genes in other appendicularian species were identified using *O. dioica* and other chordate Fgf proteins as queries for tBLASTn against publicly available genomes (i.e. *Oikopleura albicans* SCLG01000000, *Oikopleura vanhoeffeni* SCLH01000000) (Naville et al. 2019). *Fgf* genes in ascidian species other than *Ciona robusta* were identified using the Fgf protein sequences from *C. robusta* and other chordates as queries in BLASTp and tBLASTn searches against the gene models and whole genome assemblies of most species available in ANISEED (i.e. *Ciona savignyi*, *Phallusia fumigata*, *Phallusia mammillata*, *Halocynthia roretzi*, *Halocynthia aurantium*, *Botryllus schlosseri*, *Botryllus leachii*, *Molgula occulta*, *Molgula oculata* and *Molgula occidentalis*) (Brozovic et al. 2018).

Protein alignments were generated with MUSCLE and MAFFT implemented in Aliview v1.28 (Larsson 2014) and reviewed by hand. Non-homologous independently extended N- and C-terminus of different subfamilies were aligned in a non-overlapping manner to reduced background noise among Fgf subfamilies in which no similarity was detected outside the FGF core homology domain. Phylogenetic trees were based on Maximum Likelihood (ML) inferences calculated with PhyML v3.0 (Guindon et al. 2010), as well as IQ-Tree (Nguyen et al. 2015). LG was inferred as the best-fit substitution model according to Bayesian information criterion BIC to Fgf data, with a gamma with 4 categories and a shape alpha of 2.4681 (Kalyaanamoorthy et al. 2017). Tree node support was inferred by fast likelihood-based methods aLRT SH-like, aLRT chi2-based and aBayes, and by standard or ultrafast bootstraps (n=100) according to computational capacity. Phylogenetic trees were inferred both in complete and trimmed protein alignments, the later by removing all extended regions outside the FGF core homology domain, and in both cases producing the same tree topology regarding the *Fgf* subfamily homology of appendicularian genes. Gene names were assigned according to previous literature, and those that were described for the first time in this work were assigned according to the topology in the phylogenetic tree. Paralogs were named with letters in alphabetical order, reflecting orthology when the tree topology was conclusive, but not necessarily when additional species-specific duplications occurred, or the tree topology was not fully solved.

### Protein structure analyses

The domain architecture and functional motifs of Fgf proteins were examined individually with InterProScan, a comprehensive software suite that combines the information stored in several protein databases (including InterPro, PFAM, SMART or PANTHER) to provide in silico functional characterization of queried protein sequences (Jones et al. 2014). Hydropathy plots were generated in ProtScale available in Expasy (Gasteiger et al. 2005), using the Kyte-Doolittle hydrophobic scale and an interval of 9 amino acids with a linear weight variation model in a normalized scale, following previous similar analysis reported on Fgf9 (Miyakawa et al. 1999). Sequence identity and similarity for every pair of Fgf sequences was obtained from the global pairwise sequence alignment with the Needleman– Wunsch algorithm implemented in EMBOSS needle (Madeira et al. 2024). Three-dimensional structures of *O. dioica* Fgf proteins were predicted de novo with AlphaFold2 (Jumper et al. 2021). For each Fgf protein, the top-ranked relaxed model was imported into USCF ChimeraX for its visualization, analysis, and image generation (Pettersen et al. 2021). Nuclear Localization Signals (NLS) were predicted with the NLStradammus software using the 4-state HMM static model and a posterior cutoff of 0.5 (Nguyen Ba et al. 2009), or manually identified in the case of classical NLS based on the consensus stated in (Lu et al. 2021). Signal peptide (SP) predictions were conducted using the SignalP 6.0 (Teufel et al. 2022) and Phobius (Käll et al. 2004) software.

### Cloning and expression analyses

*O. dioica Fgf* genes were PCR amplified from cDNA or gDNA obtained from individuals from the Barcelona population as previously described in Martí-Solans et al. 2016. The PCR products were cloned using the Topo TA Cloning Kit (K4530-20, Invitrogen), and the resulting plasmid was digested with the adequate restriction enzyme to synthesize antisense digoxigenin (DIG) riboprobes for whole-mount in situ hybridization (WMISH) (Bassham and Postlethwait 2000; Cañestro and Postlethwait 2007; Martí-Solans et al. 2016). Primers used for the cloning of *O. dioica Fgf* genes, as well as the DNA used as template and the length of the clone and probe are indicated in **Supplementary Table 1**.

## Results

### Extensive loss of *Fgf* subfamilies in appendicularian tunicates

The gold-standard chromosome arm level genome assembly of three cryptic species of the appendicularian *O. dioica* (BAR, OSA and OKI) (Plessy et al. 2024) has allowed us to identify the full *Fgf* catalogue made up of 10 genes in each species (**Supplementary Table 2**). Phylogenetic analysis showed a clear one-to-one orthology between each *Fgf* gene in the three *O. dioica* cryptic species, providing strong evidence that we had identified the full catalogue of *Fgf* genes in *O. dioica*. Importantly, the use of three independently assembled genomes in our gene survey minimized the possibility of undetected unassembled regions containing additional *Fgf* genes, providing further evidence that the full *Fgf* catalogue in *O. dioica* was made of 10 genes. Phylogenetic analyses provided strong evidence with high node support values indicating that the 10 *Fgf* genes were paralogs originated by two bursts of appendicularian-specific duplications (red solid circled nodes in **Figure 1**). The phylogenetic tree topology also indicated with high support values that all *O.dioica Fgf* genes belonged to only two subfamilies: *Fgf11/12/13/14* and *Fgf9/16/20* (black solid circled nodes in **Figure 1**). Analyses on gene structure, protein domains and protein sequence motifs further supported this phylogenetic classification (see below). In contrast to *O. dioica*, our survey of 11 ascidian species’ genomes revealed that most of them had 7 *Fgf* genes orthologous to those previously described in *C. robusta*, which are representatives of all chordate *Fgf* subfamilies except *Fgf1/2* and *Fgf3* (Satou et al. 2002; Popovici et al. 2005; Oulion et al. 2012). The exceptions were *Ciona savingyi*, in which we could not identify an ortholog for *Fgf4/5/6*, and *Halocynthia roretzi*, *Halocynthia auriantum*, *Botrylloides leachii* and *Botryllus spp*., in which we could not identify orthologs for *Fgf-NA1*. We also surveyed the genomes of two additional appendicularians species whose genome assemblies were not too fragmented (i.e. *Oikopleura albicans* and *Oikopleura vanhoeffeni*) (Naville et al. 2019). All *Fgf* genes identified in these species belonged to the same two subfamilies found in *O. dioica* (**Figure 1** and **Supplementary table 3**). These findings across ascidian and appendicularians tunicates suggested that the loss of *Fgf1/2* and *Fgf3* likely occurred in the last common ancestor of all tunicates, before the divergence of these two groups. Interestingly, while ascidians have not systematically lost any further *Fgf* subfamily, the appendicularian ancestor lost four additional subfamilies (i.e. *Fgf4/5/6*, *Fgf7/10/22*, *Fgf8/17/18/24*, *Fgf19/21/23*) before the radiation of this clade (**Figure 1**). Regarding the evolutionary origin of *Fgf11/12/13/14*, which until now has remained unclear (Oulion et al. 2012), our phylogenetic inferences suggested that none of the cephalochordate *Fgf* genes belonged to this subfamily. This subfamily appeared to be restricted to tunicates and vertebrates (**Figure 1**), supporting previous work suggesting that *Fgf11/12/13/14* was not present in amphioxus (Bertrand et al. 2011), nor in ambulacrarian genomes, including hemichordates (Oulion et al. 2012; Fan and Su 2015) and echinoderms (Lapraz et al. 2006; Röttinger et al. 2008; Czarkwiani et al. 2021). Our results, therefore, suggested that the *Fgf11/12/13/14* subfamily was an innovation of olfactores,

**Figure 1.**
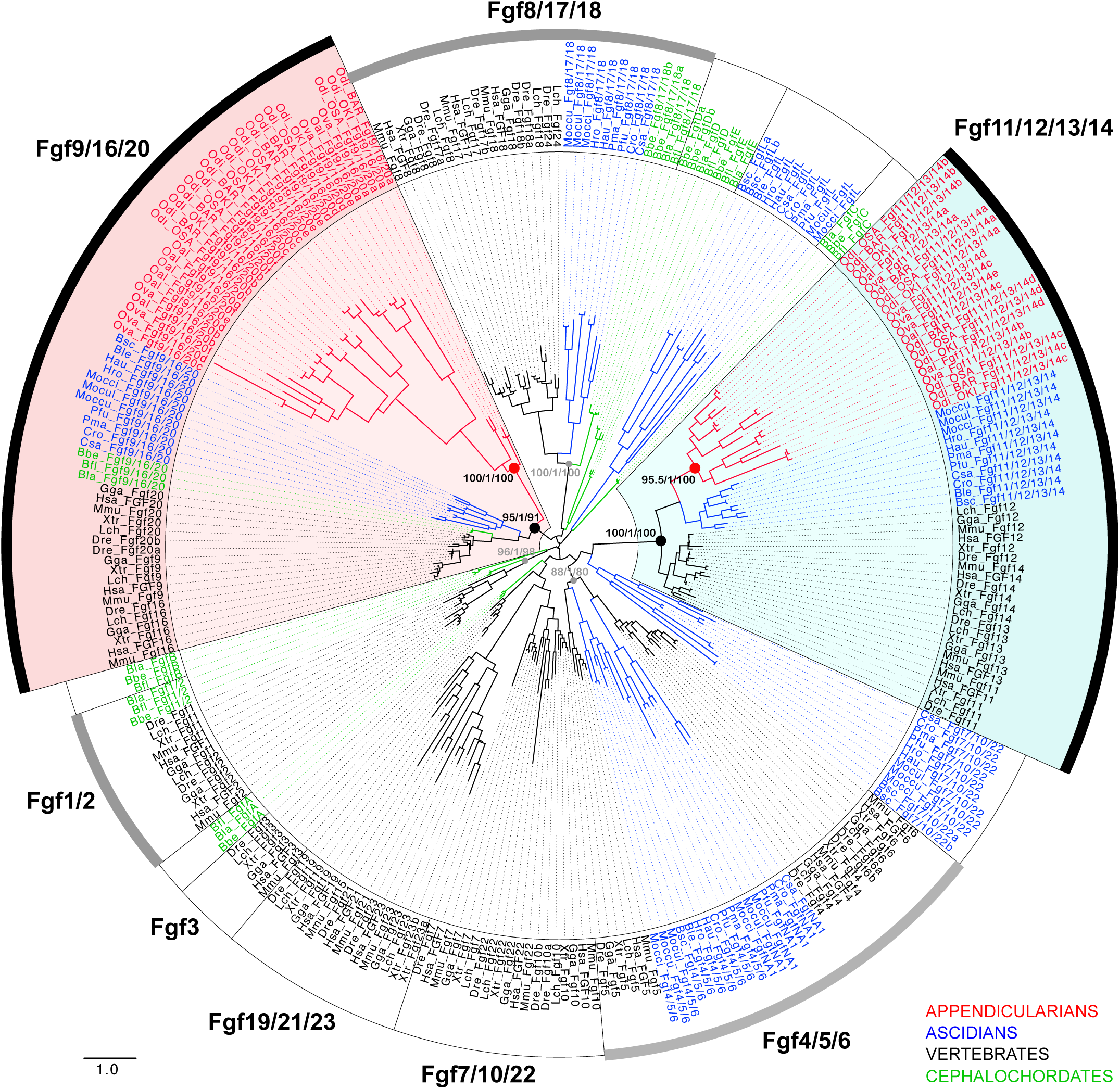
Evolutionary tree of the *Fgf* subfamilies in chordates. ML phylogenetic tree of the *Fgf* family in chordates reveals that the 10 *Fgf* genes found *Oikopleura dioica*, together with all other genes found in other appendicularians species (in red) group in two clusters with high support values (nodes with red solid circles). The tree topology indicates that the two clusters belong with high support (nodes with black solid circles) to two subfamilies, *Fgf9/16/20* (red background) and *Fgf11/12/13/14* (blue background). The presence of *Fgfs* from ascidians (in blue), vertebrates (in black) and cephalochordates (in green) allowed to infer that appendicularians has lost the subfamilies *Fgf8/17/18*, *Fgf19/21/23*, *Fgf7/10/22* and *Fgf4/5/6*. The absence of *Fgf11/12/13/14* in cephalochordates suggests that this subfamily might be a synapomorphy of the olfactores. Well supported nodes of other Fgf subfamilies (eBayes=1) with members of more than one subphylum are indicated with grey solid circles. Node support values correspond to likelihood-based method aLRT-SH-like/aBayes/uf-boostrap. The scale bar indicates amino-acid substitutions. Species abbreviations: Vertebrates (in black): *Danio rerio* (Dre), *Gallus gallus* (Gga), *Homo sapiens* (Hsa), *Latimeria chalumnae* (Lch), *Mus musculus* (Mmu), *Xenopus tropicalis* (Xtr); Ascidian tunicates (in blue): *Botrylloides leachii* (Ble), *Botrylloides schlosseri* (Bsc), *Ciona robusta* (Cro), *Ciona savignyi* (Csa), *Halocynthia aurantium* (Hau), *Halocynthia roretzi* (Hro), *Molgula occidentalis* (Mocci), *Molgula occulta* (Moccu), *Molgula oculata* (Mocul), *Phallusia fumigata* (Pfu), *Phallusia mammillata* (Pma); Appendicularian tunicates (in red): *Oikopleura albicans* (Oal), *Oikopleura dioica* (Odi), *Oikopleura vanhoeffeni* (Ova); Cephalochordates (in green): *Branchiostoma belcheri* (Bbe), *Branchiostoma floridae* (Bfl), *Branchiostoma lanceolatum* (Bla)

### Appendicularian expansion of the surviving *Fgf11/12/13/14* and *Fgf9/16/20* subfamilies

The phylogenetic analysis showed that the massive loss of *Fgf* subfamilies during the evolution of appendicularians was accompanied by a burst of duplications of the two surviving subfamilies, resulting in four *Fgf11/12/13/14a-d* paralogs and six *Fgf9/16/20a-f* paralogs (**Figure 1**). The fact that all *Fgf* genes identified in the other two analyzed appendicularians species (namely *O. vannhoeffeni* and *O. albicans*) also appeared as paralogs within these two subfamilies indicated that the expansions might predate the radiation of the clade. Moreover, the tree topology also suggested that further independent lineage-specific gene duplications might have occurred within each *Fgf* subfamily (**Figure 1**).

Analysis of microsynteny revealed a strong conservation of neighboring genes of each *Fgf* ortholog across all *O. dioica* cryptic species (i.e. Barcelona, Osaka, and Okinawa), providing strong support to the assigned homologies (**Figure 2A** and **Supplementary Figure 1**). Despite this overall microsyntenic conservation, we also observed occasional small inversions and translocations of neighboring genes in most *Fgf* gene neighborhoods. Additionally, we noted positional changes of the *Fgf* orthologs within the same chromosomal arm among the three cryptic species, consistent with the characteristic genome scrambling described in *O. dioica* (**Figure 2B** and **Supplementary Table 2**) (Plessy et al. 2024). We did not detect synteny conservation among *Fgf* paralogs within each subfamily, reinforcing the idea that most of the *Fgf* gene duplications that expanded each subfamily were not recent but likely ancestral, occurring before the radiation of the appendicularian clade.

**Figure 2.**
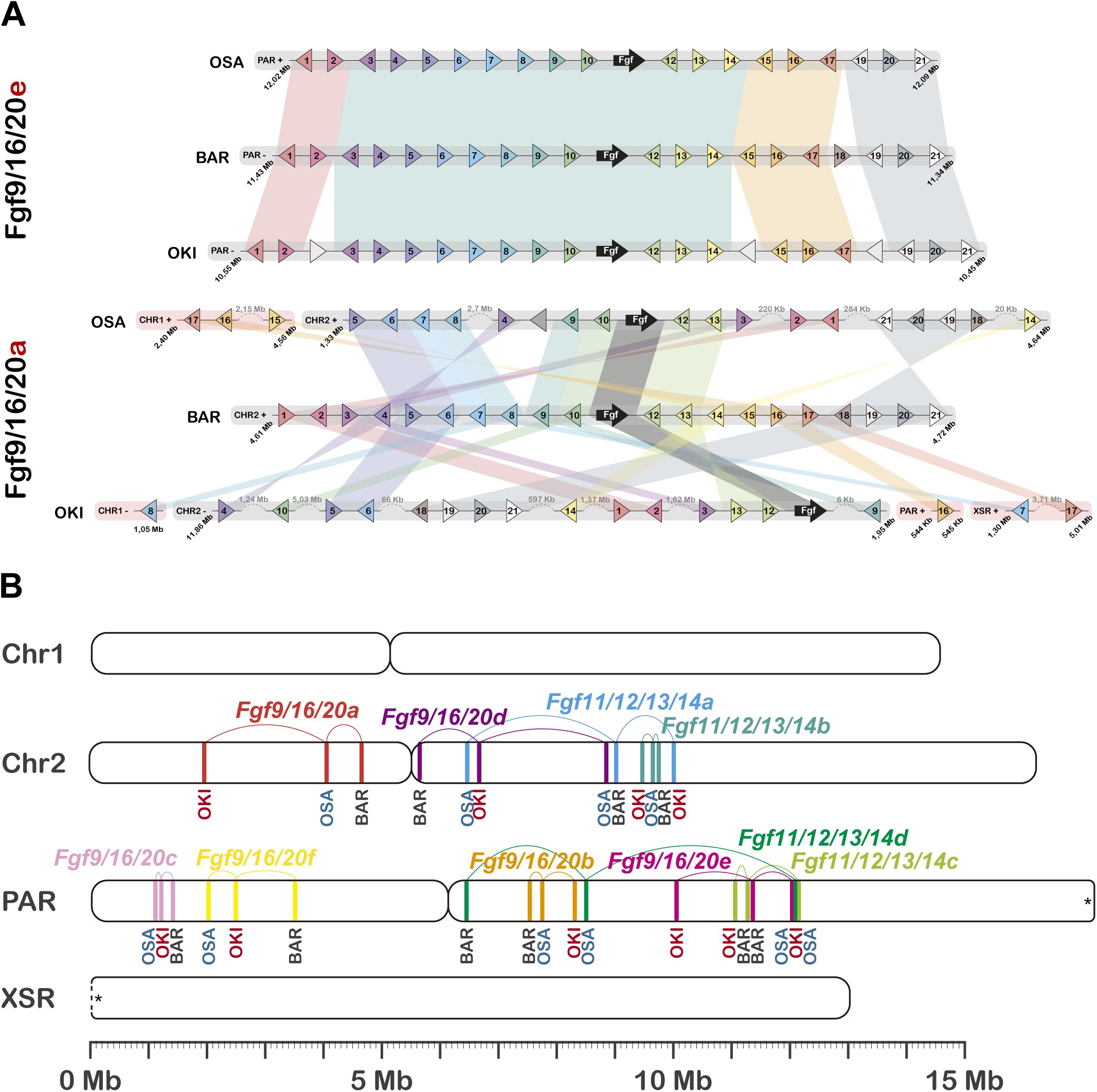
Comparative synteny analysis of *Fgf* genes among the three *Oikopleura dioica* cryptic species from Barcelona (BAR), Osaka (OSA), and Okinawa (OKI). (A) Comparison of microsynteny conservation between the genomic neighborhoods of *Fgf* genes (black arrow, and ten adjacent genes on each side). The BAR genome was used as the reference. Here, we show two illustrative examples, in which the *Fgf9/16/20e* neighborhood represents a case of high level of microsynteny conservation, and the *Fgf9/16/20a* neighborhood represents a case of low level of microsinteny conservation, especially when compared with OKI. The microsynteny comparison of the full *O. dioica Fgf* catalogue is provided in **Supplementary Figure 1**. (B) Macrosynteny analysis comparing the position of *Fgf* genes at arm chromosome level. Each *Fgf* ortholog is labeled with a distinctive color.

Comparative analysis of gene structure among appendicularian *Fgf*s and those from cephalochordates, ascidians, and vertebrates provided further support to the conclusion that all *Fgf* genes in *O. dioica* belonged to only two subfamilies. The observation that all *Fgf* genes in amphioxus retain two conserved introns in the core FGF domain (i.e. internal core intron 1 and 2: ici1 and ici2) suggested that the ancestral *Fgf* gene also had these two introns. In the *Fgf11/12/13/14* subfamily, all vertebrate members had retained these two ancestral introns, while in appendicularian and ascidian tunicates, ici1 had been lost, and only ici2 had been preserved in some of their members (**Figure 3**). The presence of an additional internal core intron (ici3) exclusively in all *O. dioica Fgf11/12/13/14* paralogs, and its identification in some *Fgf11/12/13/14* genes from other appendicularian species, further supported the notion that they were paralogs resulting appendicularians-specific duplications and suggested that ici3 could be a synapomorphic feature of the *Fgf11/12/13/14* subfamily in this lineage. Moreover, the fact that we identified two core-flanking introns (i.e. cfi1 and cfi2) in all vertebrate and tunicate *Fgf11/12/13/14* genes that were absent in all *B. floridae* and ambulacrarian *Fgf* genes, suggested that these two core-flanking introns could be a conserved synapomorphy of the *Fgf11/12/13/14* subfamily innovated in the clade olfactores (**Figure 3**).

**Figure 3.**
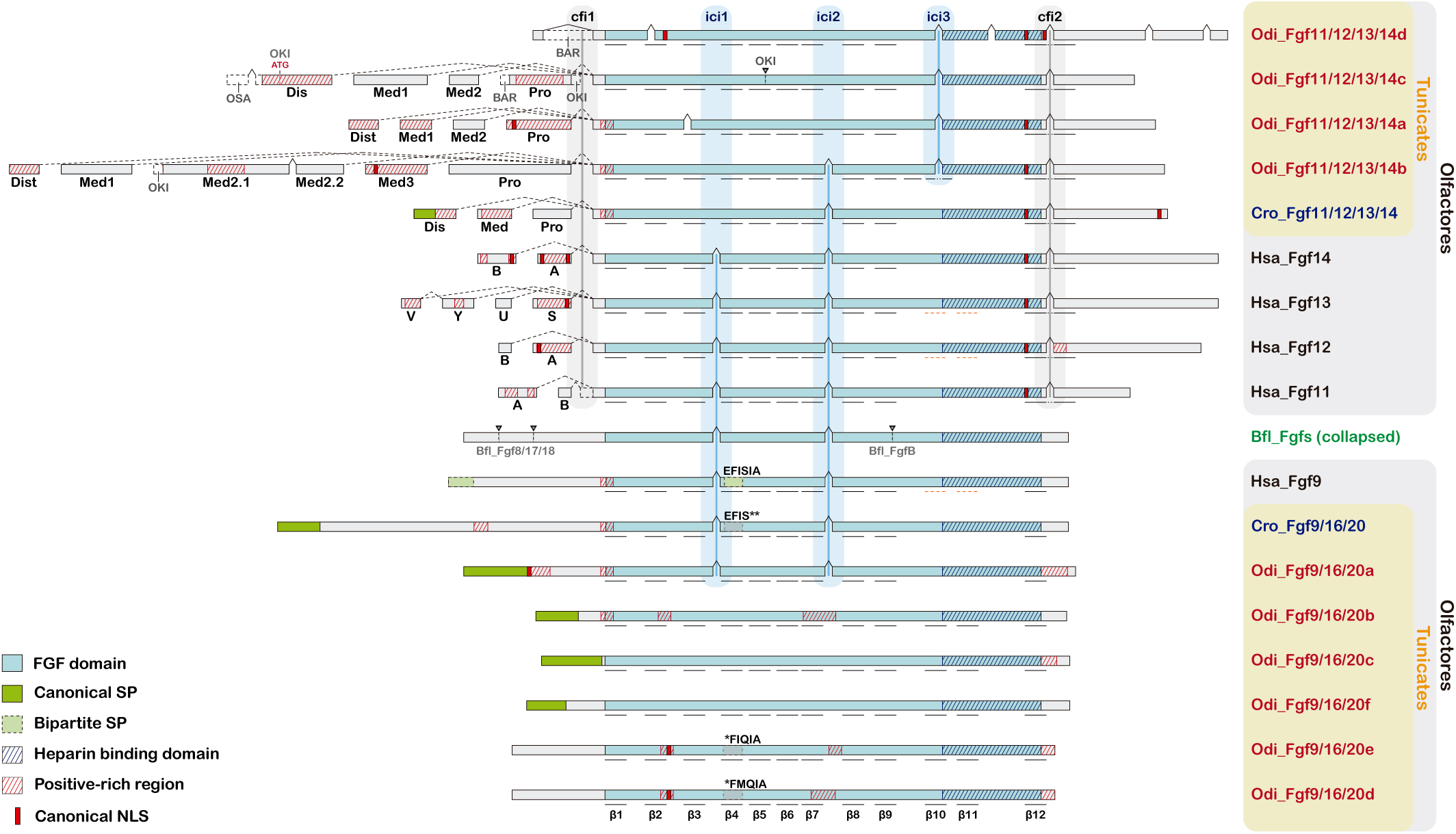
Comparative gene structures of *O. dioica* and other chordate *Fgf* genes. Exon intron organization supports phylogenetic classification of *Fgf9/16/20* and *Fgf11/12/13/14* paralogs of *O. dioica*. “cfi” denotes conserved FGF domain flanking introns, and “ici” denotes conserved internal FGF domain introns. Bfl_Fgfs represent the common structure of cephalochordate *Fgf* genes, featuring two internal core introns (ici) within the FGF domain coding sequence. Gene-specific introns are depicted as arrowheads and dashed lines at their respective locations. Predicted functional motifs are indicated as described in the legend. Black underlines highlight the presence and location of β-sheets as predicted by AlphaFold2. Orange dashed underlines highlight the presence and location of β-sheets that have been empirically determined, even though the AlphaFold2 software does not predict them (Goetz et al., 2009; Olsen et al., 2003; Plotnikov et al., 2001). For comparative purposes, genes and motifs are not drawn to scale. Dashed lines indicate alternative splicing variants of *Fgf11/12/13/14* (Dis = distal, Med = medial, Pro = proximal), and black dashed lines boxes indicate exon length differences between *O. dioica* cryptic species.

Mapping of RNAseq and EST data in the genomes of the three *O. dioica* cryptic species revealed at least four alternative first exons (namely, distal, proximal, and middle 1, middle 2, etc.) that could give rise to different isoforms due to alternative splicing and transcription start site usage in most *Fgf11/12/13/14* paralogs (**Figure 3**). The presence of alternative isoforms differing in their N-terminus in members of the *Fgf11/12/13/14* subfamily has been also described in vertebrates (Munoz-Sanjuan et al. 2000; Pablo and Pitt 2016). Our genome database surveys revealed evidence of similar alternative splice variants of the first exon in *Ciona robusta* (GeneID:445758 in NW_004190431.2) and other ascidian species, as we have also found in *O. dioica,* thus suggesting that this feature might be an ancestral characteristic of this *Fgf* subfamily in olfactores. Although many of these alternative first exons were rich in positively charged residues (i.e. Lysines and Arginines) and displayed similar hydrophobicity profiles among paralogs, sequence conservation was poor, supporting the hypothesis that the duplications that originated them were ancient in the evolution of appendicularians (**Figure 3**). The presence of small differences in the alternative splice variants among the three *O. dioica* cryptic species (e.g. the first intron of the *Fgf11/12/13/14d* has been incorporated in the open reading frame in BAR, but not in OSA or OKI), together with the presence of cryptic species-specific introns (i.e. *Fgf11/12/13/14c* showed a unique intron in the FGF domain exclusively in OKI) illustrated the rapid evolution of the *Fgf* genes in appendicularians, and provided an example of genetic variation among the cryptic species (**Figure 3**).

Analysis of gene structure of *Fgf9/16/20* paralogs in *O. dioica* revealed that while *Fgf9/16/20a* had retained the 2 ancestral introns in the FGF domain (e.g. ici1 and ici2), which were also conserved in human and ascidian *Fgf9/16/20* orthologs, all the other *O. dioica Fgf9/16/20* paralogs (i.e. *Fgf9/16/20b-f*) showed an intronless structure (**Figure 3**). This intronless structure suggested that an ancestral *Fgf9/16/20a*-like gene, most similar in sequence to *Fgf9/16/20* orthologs in other chordate species, could have been duplicated by the integration of a retrotranscribed form followed by further gene duplications. The absence of introns in most *Fgf9/16/20* genes found in other appendicularian species further reinforced the idea that this subfamily was expanded ancestrally in this clade, and that the six *Fgf9/16/20* paralogs in *O. dioica* belonged to the same subfamily. Further evidence supporting that all *O. dioica Fgf9/16/20* paralogs belonged to this subfamily was the conservation of two cysteine residues that were present in all tunicate *Fgf9/16/20* orthologs and absent in all other *Fgf* subfamilies (**Supplementary Figure 3**).

### Canonical Fgf9/16/20 paracrine and Fgf11/12/13/14 intracellular functions

Analysis of protein sequence similarity among Fgfs revealed that during the expansion of the two surviving subfamilies in appendicularians, one or two of their members (namely, Fgf9/16/20a and Fgf11/12/13/14a-b) conserved high similarity with their co-orthologs in other chordate species, while the other paralogs suffered a remarkable sequence divergence, especially within the Fgf9/16/20 subfamily. Thus, for instance, while sequence identity among paralogs of the Fgf9/16/20 subfamily in humans ranges from 62%-69.6% throughout the entire protein (80.6%-87.8% throughout the FGF core), in *O. dioica* sequence similarity among some Fgf9/16/20 paralogs was as low as 18.7% (21.2% in the FGF core) (**Supplementary table 4**). To understand how this sequence divergence might have impacted the function of the paralogs within each subfamily, we examined their conserved protein domains with HMMscan, as well as the presence of potential signal peptides (SP), nuclear localizations signals (NLS), or other conserved motifs known to interact with other cofactors and proteins.

In the Fgf9/16/20 subfamily, despite the variability of the HMMscan e-values of the FGF domains, ranging from 1^-35^ of the Fgf9/16/20a to 1^-7^-1^-13^ of the Fgf9/16/20b-f, the AlphaFold2 software predicted that most *O. dioica* Fgf9/16/20 paralogs displayed the characteristic twelve β-sheets conforming the typical β-trefoil fold structure of vertebrate Fgf9/16/20 proteins (**Figure 3** and **Supplementary Figures 2 and 3**). Moreover, the presence of regions enriched in positively charged residues (e.g. Arginine and Lysine) in all *O. dioica* Fgf9/16/20 paralogs, especially near the end of the FGF domain where heparin binding sites (HBS) have been identified in vertebrates (**Figure 3** and **Supplementary Figure 3**) (Xu et al. 2012), suggested that these regions could bind heparin or heparan sulfate proteoglycans. The presence of a β-trefoil fold and HBS, therefore, suggested that all *O. dioica* Fgf9/16/20 paralogs could potentially interact with Fgf receptors on the cell surface to function through the canonical FGF signaling. To investigate the secretion potential of *O. dioica* Fgf9/16/20 paralogs, we examined the presence of signal peptides (SP) with signalP and Phobius softwares. The results predicted the presence of SP (likelihood>0.5) in the N-terminus of four of the Fgf9/16/20 paralogs (namely, Fgf9/16/20abc and f), but not in the other two (d and e). The presence of SPs in four of the Fgf9/16/20 paralogs was comparable to the SP described in the N-terminus of Fgf9/16/20 of *C. robusta* (Satou et al. 2002), and it can explain the degeneration of the non-canonical signal peptide EFISIA motif within the FGF core, which is required for extracellular secretion via the endoplasmic reticulum in other organisms (Popovici et al. 2004). Interestingly, the absence of an SP in Fgf9/16/20d-e correlated with a certain conservation of the EFISIA motif (i.e. TFIQIA), producing a peak of hydrophobicity like those observed in the EFISIA motif of Fgf9/16/20 in other species (**Figure 3** and **Supplementary Figure 4**) (Miyakawa et al. 1999; Popovici et al. 2004). These observations, therefore, further supported the paracrine nature of the Fgf9/16/20 subfamily in appendicularians and suggested that different paralogs might have evolved different secretion mechanisms.

In the Fgf11/12/13/14 subfamily, its members have been traditionally associated with intracellular functions through interacting with various proteins. such as regulators of voltage-gated channels (i.e. Na_v_s or Ca_v_s), as regulators of transcription factors (i.e. islet brain-2 or NEMO), or as players of neuronal cytoskeleton architecture and cell morphology (Pablo and Pitt 2016). Like their vertebrate counterparts, all appendicularian Fgf11/12/13/14 paralogs lacked a signal peptide (**Figure 3**), providing the first clue of a conserved intracellular function. The conservation in all *O. dioica* Fgf11/12/13/14 paralogs of a Leucine and an Arginine in positions that have been described to be critical for the interaction with Na_v_s and islet brain-2 (Olsen et al. 2003; Pablo and Pitt 2016), and that were conserved in all vertebrate and ascidian members of the Fgf11/12/13/14 subfamily, but not in other Fgf subfamilies, reinforced the idea that this subfamily also played an intracellular function in appendicularians (**Supplementary Figure 3**).

The presence of multiple alternative splice variants with different first exons found in most *O. dioica Fgf11/12/13/14* paralogs (**Figure 3**) was also consistent with the same feature described in vertebrates for *Fgf11/12/13/14* genes with intracellular functions related to the modulation of voltage-gated channels regulating neural excitability (Munoz-Sanjuan et al. 2000; Laezza et al. 2009).

Upon the growing evidence that Fgf proteins may also have intranuclear functions (Popovici et al. 2006), we conducted NLS predictions with NLStradamus and searched for KRVR motifs known to provide NLS in *O. dioica* (Clarke et al. 2007). Our analysis revealed that most *O. dioica* Fgf11/12/13/14 paralogs possess an NLS at the end of the FGF core, similarly to vertebrate and ascidian Fgf11/12/13/14 proteins. Interestingly, we also found NLS in some of the alternative first exons that generate different isoforms diverging at the N-terminus, suggesting that these different isoforms not only might have different promoter usage, but also different intracellular localizations (**Figure 3**). We also found NLS in three out of the six *O. dioica* Fgf9/16/20 paralogs, including the two that lack an SP, implying that these paralogs may have also evolved non-secreted functions.

Overall, our findings from the structural analysis were consistent with paracrine functions for the *Fgf9/16/20* subfamily and intracellular functions for the *Fgf11/12/13/14* subfamily in appendicularians. The high sequence divergence, variation in the presence of putative SP and NLS, and the formation of different isoforms due to differential splice variants in the N-terminus raised the possibility that multiple functions might have also evolved among the different paralogs duplicated during the expansion of these two surviving families in appendicularians.

### *Fgf* expression atlas during the development of *Oikopleura dioica*

To better understand the functional consequences of gene loss and gene expansion on the *Fgf* subfamilies in *O. dioica*, we performed an exhaustive expression analysis of all the *Fgf9/16/20* and *Fgf11/12/13/14* paralogs by whole mount in situ hybridization (WISH) throughout development, from eggs to late-hatchling stages (**Figure 4 A-J**). In general, we found that the level of expression signal of most *Fgf* genes was low, and long periods of staining (e.g., between 1 to 14 days) were required to visualize some of the tissue-specific expression domains. Our description here will mostly focus on tissue-specific *Fgf* expression domains repeatedly observed in different embryos over background levels, but we cannot discard that in addition to those specific domains some of the *Fgf* genes also had a generalized basal expression scattered in other parts of the embryo, as it has been also described in other animals including *C. robusta* and amphioxus (Imai et al. 2004; Bertrand et al. 2011).

**Figure 4.**
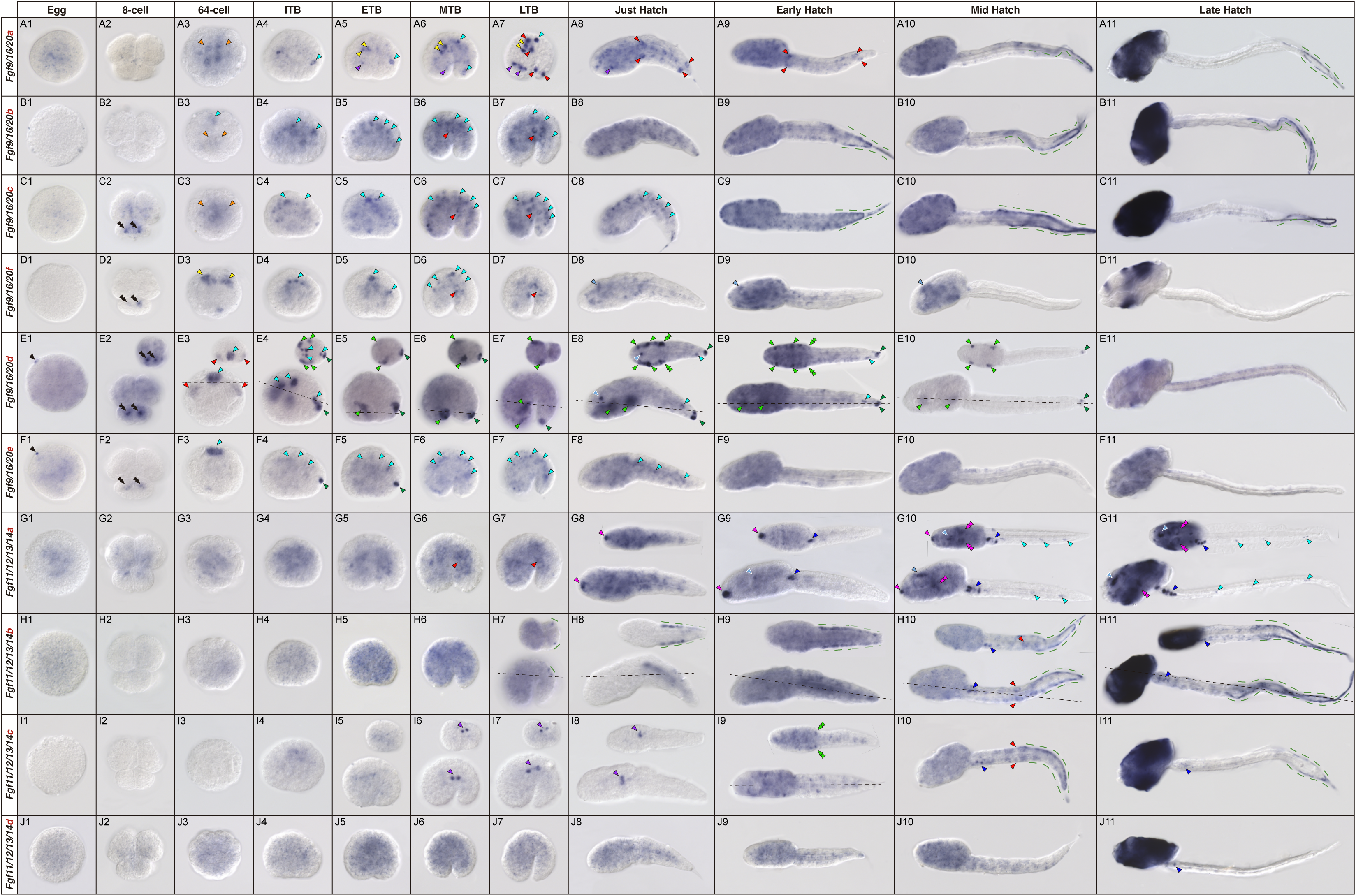
Developmental expression atlas of *O. dioica Fgf* genes. Whole-mount in situ hybridization images of *O. dioica* at various developmental stages: eggs (A-J1), 8-cell embryos (A-J2), 64-cell embryos (A-J3), incipient tailbud (ITB) embryos (A-J4), early tailbud (ETB) embryos (A-J5), mid tailbud (MTB) embryos (A-J6), late tailbud (LTB) embryos (A-J7), just hatchlings (A-J8), early hatchling larvae (A-J9), mid hatchling larvae (A-J10), and late hatchling larvae (A-J11). Central images in each panel are left lateral views, oriented anterior to the left and dorsal to the top. Upper-right image insets (’) are dorsal views of optical cross-sections at the levels indicated by black dashed lines. Black arrowheads label an stained cortical spot in unfertilized eggs; black double arrowheads label the A pair blastomeres in 8-cell embryos; orange arrowheads mark ingressing vegetal blastomeres in the 64-cell embryos; blue arrowheads label neural derivatives (cyan-blue labels the neural plate in 64-cell and ITB embryos, and the nerve cord in later stages; dark-blue labels the caudal ganglion, and pale-blue labels the anterior brain); light green arrowheads label epidermal domains in the trunk and light green double arrowheads label the area of the Langerhans receptors primordia; dark green arrowheads label epidermal domains in the tailbud tip; green dashed lines mark the lateral epithelium of the tail and the fins; purple arrowheads label undetermined endomesodermal domains in the trunk; magenta arrowheads mark the mouth primordium; magenta double arrowheads mark the pharyngeal slits; yellow arrowheads label notochord cells; red arrowheads label muscle precursor cells and muscle cells in the tail.

In oocytes, some *Fgf* genes (i.e. *Fgf9/16/20a,d,e*, and *Fgf11/12/13/14a,b,d*) showed a weak staining signal in the cytoplasm that was difficult to distinguish over the staining background. This suggested that, in general, *Fgf* transcripts were not a major component of the maternal contribution (**Figure 4 A-J1**). In the case of *Fgf9/16/20d* and *Fgf9/16/20e*, however, we observed a small but intense staining spot in the cortical area of many unfertilized eggs (n=11/13 and n=9/11, for *Fgf9/16/20d* and *Fgf9/16/20e*, respectively), but not in all. This suggested that the formation of this *Fgf* transcripts spot might be transient and difficult to capture, or perhaps not present in all individuals (**Figure 4 E-F1, black arrowheads**). In the early stages of development, from 8-to 64-cell, all *Fgf9/16/20* paralogs started showing staining signals, suggesting that their expression onset took place concomitant with the activation of zygotic transcription (Wang et al. 2015) (**Figure 4 A-F2 & A-F3**). At the 8-cell stage, many *Fgf9/16/20* paralogs (i.e. *Fgf9/16/20c*, *Fgf9/16/20d*, *Fgf9/16/20e* and *Fgf9/16/20f*) showed staining signal restricted to the smaller pair of blastomeres in the vegetal pole (**Figure 4 C-F2, black double arrowheads**). These cells corresponded to the A/A4.1 blastomere pair in Delsman/Conklin nomenclature, which gives rise to most of the nervous system, the notochord, and other endomesodermal derivatives (Nishida 2008; Stach et al. 2008). At the gastrula stage (32-64 cells), we found that all *Fgf9/16/20* paralogs were expressed in the precursor blastomeres of either mesodermal or ectodermal derivatives. *Fgf9/16/20a*, *Fgf9/16/20b* and *Fgf9/16/20c* staining was detected in the endomesodermal blastomeres (**Figure 4 A-C3, orange arrowheads**); *Fgf9/16/20b*, *Fgf9/16/20d* and *Fgf9/16/20e* staining was detected in the neural plate (**Figure 4 B3 & E-F3, cyan arrowheads**); *Fgf9/16/20f* was detected in notochord precursor cells (**Figure 4 D3, yellow arrowheads**); and *Fgf9/16/20d* staining was detected in muscle precursor blastomeres (**Figure 4 E3, red arrowheads**). Consistent with these observations in early developmental stages, the majority of tissue-specific *Fgf* expression domains observed during later embryogenesis were also predominantly associated with ectodermal derivatives (e.g., nervous system and epidermis) or mesodermal derivatives (e.g., notochord and muscle).

Among ectodermal derivatives, the staining signal of *Fgf9/16/20* paralogs observed in the neural plate of 64-cells stage embryos persisted in cells of the developing nervous system up to the hatchling stage. These signals were observed in precursors of the brain, the caudal ganglion, and the spinal cord (**Figure 4 A-F**, **different tones of blue arrowheads**). The fact that some of these neural expression domains disappeared at specific developmental stages suggested that some *Fgf9/16/20* paralogs might be precisely regulated in specific subsets of neural populations along the anteroposterior axis during the formation of the central nervous system. For instance, *Fgf9/16/20d* expression domains were clearly observed in the neural plate at 64-cell stage and in the posterior part of the brain in incipient tailbud (ITB) embryos (**Figure 4 E3-4, cyan arrowheads**), but it was absent in subsequent stages until the just-hatchling stage, when it reappeared in the developing brain (**Figure 4 E8, light blue arrowhead**). We also observed staining signals for *Fgf11/12/13/14* paralogs in neural domains. However, in contrast to *Fgf9/16/20* genes, which were predominantly expressed before hatching, *Fgf11/12/13/14* genes were mainly expressed during the hatchling stages (**Figure 4 G-J**, **different tones of blue arrowheads**). Neural expression domains of *Fgf11/12/13/14* genes were detected in various locations within the nervous system, including specific cells of the brain dorsal to the sensory vesicle, the ventral region of the ciliary funnel (**Figure 4 G10-11, light blue arrowheads**), groups of cells in the caudal ganglion (**Figure 4 G9-11, H10-11, I10-11 & J11, dark blue arrowheads**), and isolated cells at different positions along the nerve cord in the tail (**Figure 4 G10-11, cyan arrowheads**). Overall, these results revealed a complex pattern of *Fgf* expression in the developing neural system, suggesting that despite the extensive loss of *Fgf* subfamilies, the expansion of the surviving *Fgf* genes has allowed the preservation of neural functions during appendicularian development, similar to other chordates.

Among ectodermal derivatives, we also observed specific expression domains for various *Fgf* genes in the epidermis, both in the tail and in the trunk (**Figure 4**, **different tones of green arrowheads and dashed lines**). In the tail, we observed a dynamic expression pattern with different *Fgf* genes expressed at different levels of the anteroposterior axis, mainly in two areas: the developing fin and the tip of the tail. In the precursor cells of the fin, *Fgf11/12/13/14b* expression was first detected in a bilateral pair of epidermal cells located in the middle region of the tail (**Figure 4 H7, green dashed lines**). This expression later spread to the first third of the tail at the just-hatch stage and eventually extended, along with other *Fgf* paralogs (namely, *Fgf11/12/13/14c*, *Fgf9/16/20a*, *Fgf9/16/20b* and *Fgf9/16/20c*) to the posterior half of the tail in mid- and late-hatchlings (**Figure 4 A10-11, B-C9-11, H7-11 & I10-11, green dashed lines**). In the tip of the tail, a pair of epidermal cells started showing strong staining for *Fgf9/16/20d* and *Fgf9/16/20e* at the ITB stage. While *Fgf9/16/20e* expression was downregulated by the mid-tailbud stage, *Fgf9/16/20d* signal persisted until the mid-hatchling stage (**Figure 4 E4-10 & F4-5, dark green arrowheads**). In the trunk, bilateral groups of epidermal cells expressed *Fgf9/16/20d* at different levels of the anteroposterior axis (**Figure 4 E4-10, light green arrowheads**). The most posterior group included the primordia of the Langerhans receptors, which also expressed *Fgf11/12/13/14c* (**Figure 4 E8-9 & I9, light green double arrowheads**). In the most rostral region of the trunk epidermis, *Fgf11/12/13/14a* showed an expression domain in a group of subepidermal cells in the area of the mouth at the just-hatchling stage. This expression domain was later expanded to the epidermal surface by the mid-hatched stage, coinciding with the opening of the mouth (**Figure 4 G8-10, magenta arrowheads**). Interestingly, *Fgf11/12/13/14a* expression was also observed in the pharyngeal slits in the mid- and late-hatchling larvae (**Figure 4 G10-11, magenta double arrowheads**).The expression of *Fgf11/12/13/14a* in the primordia of organs in which ciliated sensory cells developed (i.e. the stomodeum, the ciliary funnel, and the ciliary rings), together with the expression of *Fgf9/16/20d* and *Fgf11/12/13/14c* in the primordia of the Langerhans receptors, suggested that FGF signaling might be involved in the development of placodial derivatives in appendicularians, as well as in structures in which epithelial perforation and fusions occur, as described in other chordates (Bassham and Postlethwait 2005; Kourakis and Smith 2007; Bassham et al. 2008). In late hatchling stages, nearly all *Fgf* genes were strongly expressed in different parts of the oikoblast, the organ responsible for the architecture and secretion of the house (**Figure 4 A-J11**). Some showed generalized patterns, while others were restricted to or excluded from specific regions. For example, *Fgf9/16/20f* was restricted to cells adjacent to the anterior cells of the field of Fol, the anterior rosette, the field of Martini, the posterior rosette, and its adjacent lateral bands (**Figure 4 D11**). In contrast, *Fgf9/16/20a-b* were expressed throughout the entire oikoblast but excluded from the ring of the mouth and the Giant cells (**Figure 4 A-B11**). This finding suggested that FGF signaling has been recruited for the development of this innovative organ responsible for the formation of the house, as described for many other developmental genes in appendicularians (Mikhaleva et al. 2018).

Among endomesodermal derivatives, the notochord exhibited an *Fgf9/16/20a* expression domain restricted to the first and third cells from the early tailbud (ETB) to late tailbud (LTB) stages (**Figure 4 A5-7, yellow arrowheads**). In these stages, *Fgf9/16/20a* staining was also observed in few internal cells bilaterally located in the anterior half of the trunk, whose positions were compatible with endomesodermal progenitors of the pharynx, endostyle or buccal glands (**Figure 4 A5-8, purple arrowheads**). At the LTB stage, a new mesodermal expression domain of *Fgf9/16/20a* appeared restricted to the first and eighth pairs of muscle cells, and it was maintained until the early hatchling stage (**Figure 4 A7-9, red arrowheads**). Other *Fgf* paralogs with broad, ubiquitous expression patterns also showed stronger expression in muscle cells compared to other parts of the embryo (i.e. *Fgf9/16/20b*, *Fgf9/16/20c*, *Fgf11/12/13/14b* and *Fgf11/12/13/14c*; **Figure 4**, **red arrowheads**). From ITB to LTB stages, two cells located on the right side of the anterior part of the notochord, which later at the early hatchling stage were located anteriorly in a rostral position to the notochord, expressed a very distinct expression of *Fgf11/12/13/14c* (**Figure 4 I6-8, purple arrowheads**). We could not determine the identity of these endomesodermal cells but, considering their position, they could be related to the development of endodermal progenitors of the digestive system or the gonad (Olsen et al. 2018).

## Discussion

### Less, but more: massive gene losses accompanied by bursts of duplications of the surviving paralogs

The study of FGF signaling is central for understanding many fundamental functions of cell biology, including proliferation, differentiation, migration, apoptosis, and survival of cells, from embryo development to adult tissue homeostasis, as well as for understanding how its malfunctioning can cause several diseases (reviewed in Dorey and Amaya 2010; Xie et al. 2020). FGF signaling has an ancient evolutionary origin, at least already present in the ancestral eumetazoan (Bertrand et al. 2014). Some of the conserved core functions of the FGF signaling also have an ancestral origin, such as mesodermal induction which predates the Cambrian explosion and the origins of Bilateria (Matus et al. 2007). Different taxa, however, have innovated a great variety of other FGF functions, in many cases associated to gene duplications, and often accompanied by gene losses (Popovici et al. 2005; Oulion et al. 2012). During the evolution of chordates, for instance, the expansion of eight single-gene subfamilies up to 27 *Fgf* genes early in the evolution of vertebrates due to the two rounds of genome duplication has been linked to the innovation and sophistication of many of the characteristic vertebrate traits, including axial patterning, somitogenesis, limb bud formation, visceral and skeletal development (Thisse and Thisse 2005), and even the invention of a “new head” (Bertrand et al. 2011). Studies of FGF signaling in ascidian tunicates have contributed to reveal that some of the traits that characterize vertebrates, indeed, were not vertebrate innovations, but were already present in the last common ancestor of olfactores, such as the role of *Fgf8/17/18* in the organizer activity and the compartmentalization of the central nervous system and in placodal derivatives (Kourakis and Smith 2007; Imai et al. 2009; Wagner and Levine 2012; Stolfi et al. 2015; Horie et al. 2018). Here, our results in *O. dioica* unveil an unprecedented case among chordates, in which massive gene losses have erased all *Fgf* subfamilies but two, *Fgf9/16/20* and *Fgf11/12/13/14*. Interestingly, the massive gene losses have been accompanied by two bursts of duplications that have given rise to several *Fgf* paralogous genes within each subfamily describing an evolutionary scenario of “less, but more” (**Figure 5A**).

**Figure 5.**
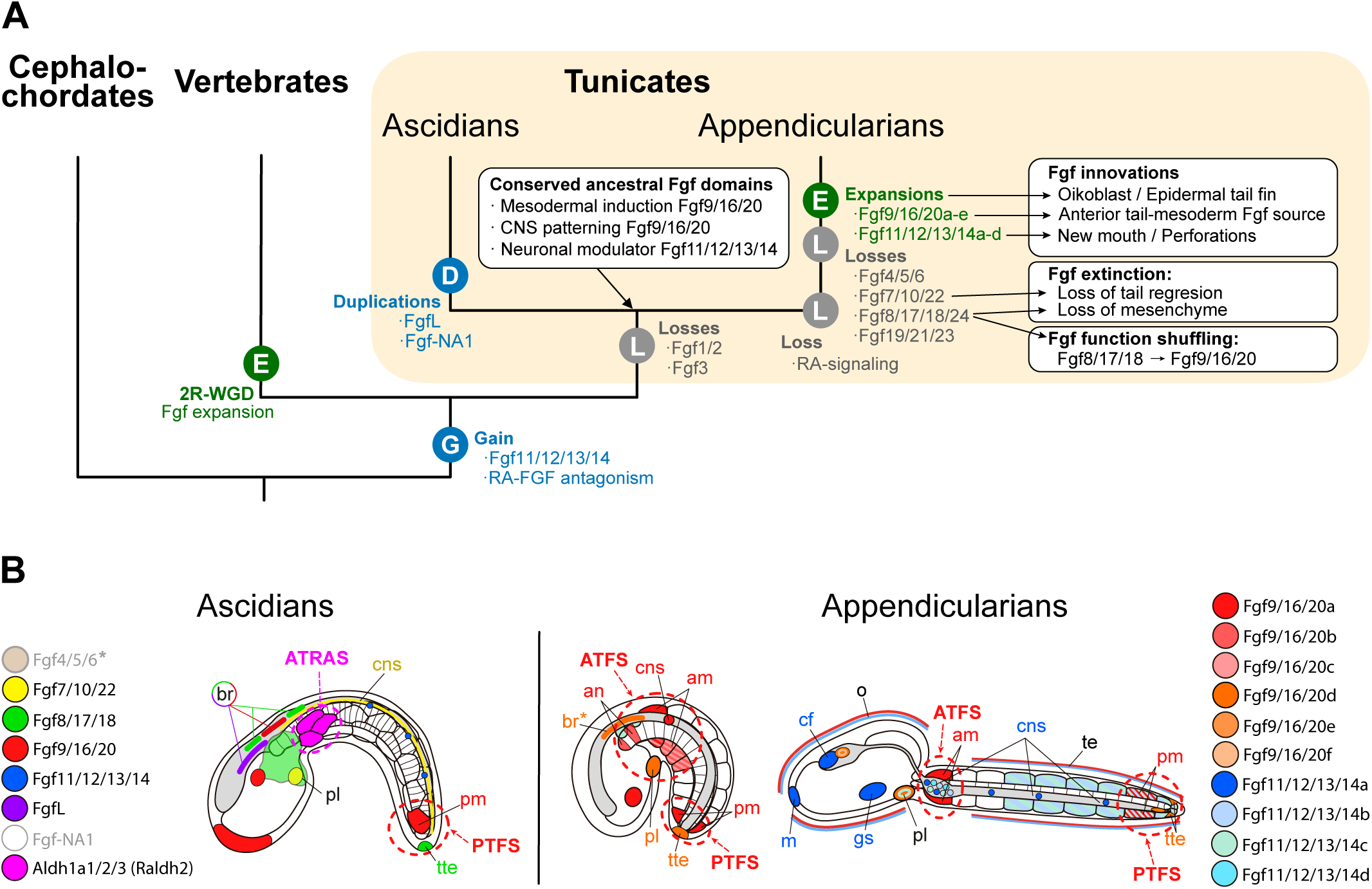
Evolutionary scenario of *Fgf* subfamilies in chordates. (A) Evolutionary tree of chordate subphyla indicating main events of losses (L), gains (G), duplications (D) or expansions by burst of duplications (E), as well as main associated patterns of conservation, innovation, function shuffling and extinction of *Fgf* expression domains. 2R-WGD: two rounds of whole genome duplication. (B) Comparative schematic representation of the main expression domains of *Fgf* subfamily members between ascidians and appendicularians (tailbud stage in the left, and hatchling in the right). Distinct colors are assigned to each *Fgf* as indicated in the figure legend. *FgfNA1* expression has not been described in ascidians, and *Fgf4/5/6*\* has been detected maternally and widely throughout development with no obvious tissue-specific domains (Imai et al., 2004). In appendicularians, expression of *Fgf9/16/20* paralogs is the most abundant in tailbud stages, and expression of *Fgf11/12/13/14* paralogs is more obvious at hatchling stages. Asterisk (as in br*) denotes that expression was found in a slightly earlier stage than the one represented in the figure. Ascidians show a characteristic PTFS (Posterior-Tail FGF Source) and ATRAS (Anterior-Tail RA Source), while in appendicularians the loss of RA signaling (Cañestro & Postlethwait, 2007; Martí-Solans et al., 2016) might be related to the innovation of an ATFS (Anterior-Tail FGF Source), considering that the conserved RA-FGF antagonistic action emerged at the base of olfactores (Pasini et al. 2012; Bertrand et al. 2015). Abbreviations: an: anterior notochord, am: anterior muscle, br: brain, cf: ciliary funnel, cns: central nervous system, gs: gill slit, m: mouth, o: oikoblastic epithilium, pl: placode, pm: posterior muscle, te: tail epidermis, tte: terminal tail epidermis.

Our analysis of gene phylogenies, synteny conservation, gene architecture and structural protein motifs across various appendicularians species provide solid evidence that all *Fgf* genes in appendicularians belong to only two subfamilies. This indicates that the losses likely occurred at the base of this clade, and that different species might have independently expanded their *Fgf* catalogue in a very dynamic fashion. This dynamic evolution of *Fgf* genes is particularly evident when comparing different cryptic species of *O. dioica*, where divergences are observed in the presence of different isoforms due to alternative splicing, significant sequence divergence outside the FGF domain, presence of novel introns and microsyntenic rearrangements. The vast conservation of the FGF domain with their typical β-trefoil topology supports the idea that Fgf paralogs in *O. dioica*, despite their sequence divergence, can function as the typical Fgf ligands of other animals. The common presence of secretion motifs in *Fgf9/16/20* paralogs suggests that members of this subfamily can act extracellularly through the canonical signaling pathway typical of this subfamily (Itoh and Ornitz 2011; Ornitz and Itoh 2015). On the other hand, the extensive presence of nuclear localization signals in Fgf11/12/13/14 paralogs suggests that members of this subfamily might have intracellular functions, as it has been described for members of this *Fgf* subfamily in other chordates (Smallwood et al. 1996; Pablo and Pitt 2016).

### Evolutionary patterns associated to the “less, but more” scenario of *Fgf* evolution in tunicates

Our comparative *Fgf* expression analysis between appendicularians, ascidians and other chordates allow us to identify cases that can be categorized into four different evolutionary patterns associated to the “less, but more” scenario (**Figure 5A**):

#### (1) conservation of ancestral expression domains

In the first category, we found that the early expression domains of the single ascidian co-ortholog *Fgf9/16/20* in the A4.1 derived vegetal blastomeres of *C. intestinalis* have been preserved in four of the *Fgf9/16/20* paralogs of *O. dioica* (i.e. *Fgf9/16/20c*, *d*, *e* and *f*) in the same equivalent vegetal blastomeres (Kaoru S. Imai et al. 2002; Bertrand et al. 2003; Hudson et al. 2016; Satou 2020). This expression has been related to an ancestral function of the FGF signaling conserved among most bilaterians in initiating mesodermal and endodermal gene regulatory networks (Technau and Scholz 2003). In ascidians, *Fgf9/16/20* is sufficient for the first phase of initiation of mesodermal induction, although *Fgf8/17/18* is also required for a second phase of fate maintenance (Yasuo and Hudson 2007). This second phase might have been modified in *O. dioica* upon the loss of the Fgf8/17/18 subfamily. Moreover, the early expression of *Fgf9/16/20* paralogs at the eight-cell stage in *O. dioica* is compatible with recent findings in ascidians in which FGF acts as a timer for zygotic genome activation whose responsiveness sharply starts between the 8- and 16-cell stage (Treen et al. 2023).

A second example of conservation of ancestral expression domains is shown by *Fgf9/16/20d-e* in the developing central nervous system from the 64-cell to tailbud stages, comparable to the expression of the ascidian *Fgf9/16/20* co-ortholog in equivalent positions in the central nervous system dorsally at the level of the anterior tip of the notochord. This similarity suggests conserved *Fgf9/16/20* ancestral functions in the anteroposterior patterning of the central nervous system among tunicates (Karou S. Imai et al. 2002; Miyazaki et al. 2007) (**Figure 5B**).

For the *Fgf11/12/13/14* subfamily, our data show that many of the expression domains of its paralogs in *O. dioica* are associated to the nervous system (i.e. dorsal and anterior ventral part of the brain, subset of cells of the caudal ganglion, as well as in isolated neurons located at different levels) (**Figure 5B**). This neural *Fgf* expression is a shared characteristic with their vertebrate homologs, many of which are expressed in neurons where they perform Fgf receptor-independent intracellular functions. These functions involve interactions with voltage-gated sodium channels, affecting neuronal excitability, and have been implicated in human neuronal diseases (Wang et al. 2011). Supporting this neuronal function, our findings show that most *O. dioica Fgf11/12/13/14* paralogs produce different isoforms corresponding to alternative splice variants of the first exons, a feature also shared with vertebrate *Fgf11/12/13/14* genes that interact with voltage-gated sodium channels (Laezza et al. 2009). In ascidians, although *Fgf11/12/13/14* has no detectable expression during development (Satou et al. 2002; Treen et al. 2014), our finding of alternative splicing in the first exon also suggests a conserved neural function. This is consistent with recent single cell RNA-seq data in ascidians showing abundant *Fgf11/12/13/14* transcripts in specific neurons of the tail, including bipolar tail neurons, which have been proposed to share properties with neural-crest-derived dorsal root ganglia (Stolfi et al. 2015; Horie et al. 2018). Future expression analyses of the ascidian *Fgf11/12/13/14* in post-metamorphic animals to uncover more types of neurons that express this intracrine Fgf ligand, as well as functional characterization of its neural role in modulating neuronal excitability, will be of great interest to develop new animal models to better understand the molecular basis of related human neuronal disorders. Moreover, considering that our results reinforce the hypothesis that amphioxus and ambulacrarians lack *Fgf11/12/13/14* (Lapraz et al. 2006; Röttinger et al. 2008; Bertrand et al. 2011; Fan and Su 2015; Czarkwiani et al. 2021), and suggest that its origin could be an innovative synapomorphy of olfactores (**Figure 5A**), further studies of this gene subfamily in appendicularians and other tunicates could shed light on the role of intracrine Fgf ligands in the evolution of the nervous system within this clade following its divergence from cephalochordates.

#### (2) function shuffling among surviving paralogs upon the loss of genes

In this second category, our study reveals two paradigmatic examples of function shuffling between the ascidian *Fgf8/17/18* or *Fgf7/10/22* and the *O. dioica Fgf9/16/20d*, all of which belong to the group of paracrine/autocrine secreted Fgf ligands. First, the *Fgf8/17/18* epidermal expression domain in the tip of the tail of ascidians, which acts as a secreted posterior tail FGF source (PTFS) (Pasini et al. 2012; Kim et al. 2020), is comparable to the equivalent domain of *Fgf9/16/20d* in the posterior tip of the tail in *O. dioica* (**Figure 5B**). Second, *Fgf8/17/18* and *Fgf7/10/22* in ascidians are also expressed during the development of the atrial siphon, which has been related to the evolution of otic placode homologs (Kourakis &Smith, 2007). In *O. dioica*, in the absence of these two genes, it is *Fgf9/16/20d* and *Fgf11/12/13/14c* the paralogs that show equivalent expression domains in the Langerhans receptor primordia, which have been proposed to be homologous placodial structures in appendicularians (Bassham and Postlethwait 2005) (**Figure 5B**).

#### (3) innovation of novel expression domains in novel paralogs

In the third category, among the novel expression domains innovated by the duplicated paralogs in *O. dioica*, we find at least three examples. First, in early hatchling stages *Fgf11/12/13/14a* is specifically expressed in the stomodeum, coinciding with the time when the mouth is opening (**Figure 5B**). No comparable expression has been observed for the ascidian *Fgf11/12/13/14* gene, which shows no detectable expression during development (Satou et al. 2002; Treen et al. 2014). Interestingly, the stomodeum and mouth opening derive from the anterior neuropore in ascidians (Veeman et al. 2010), while in *O. dioica* these structures develop in the most rostral part of the trunk directly connecting to the pharynx. In *O. dioica*, the recruitment of *Pax2/5/8a* expression in the primordium of the stomodeum has been suggested to be related to cellular functions of perforation, adhesion and fusion of epithelial openings, including the mouth (Bassham et al. 2008). Therefore, despite the expression of placodial markers such as *Pitx* suggests deep genetic homology among mouths and adenohypophysis-like organs –i.e. the ciliary funnels in ascidians and appendicularians, and the Hatschek’s pit in amphioxus– (Bassham and Postlethwait 2005) we cannot discard the possibility that the mouth of ascidians and appendicularians have independent evolutionary origins recruiting a common cassette of placodial genes as has been suggested for other placodial-derived structures (Bassham and Postlethwait 2005). The involvement of different Fgf ligands in the late development of the mouth and pharyngeal slits in various animals across diverse taxa suggests that the FGF signaling might have been repeatedly recruited during the evolution of perforated structures (Crump et al. 2004; Röttinger et al. 2008; Bertrand et al. 2011; Fan et al. 2018; Rees et al. 2024). Moreover, the high expression of *Fgf11/12/13/14a* in the stomodeum, pharyngeal slits, and in the rostro-ventral part of the brain related to the ciliary funnel, regions also expressing *Pax2/5/8a*, suggests that *O. dioica* may have innovated the recruitment of *Fgf11/12/13/14a* in the evolution of mechanisms related to ciliary cells and perforated structures. To our knowledge, this would be the first case in which an intracrine Fgf ligand has been related with the development of the mouth.

The second example is related to *Fgf* expression in mesodermal derivatives, such as the notochord or muscle cells, in the anterior region of the tail during tailbud and hatchling stages. In ascidians, no *Fgf* expression has been detected in cells of the notochord or tail muscles in tailbud/hatchling stages, except for *Fgf9/16/20* being expressed in the most posterior pair of muscle cells near the tip of the tail (Imai et al. 2004; Pasini et al. 2012). In contrast, *O. dioica* exhibits strong expression of *Fgf9/16/20a* in the most posterior muscle cells of the tail and the most anterior pair, as well as in the first and third cells of the notochord. Additionally, *Fgf9/16/20b-d* expression is observed above background levels in the three most anterior and central muscle cells at tailbud stages. These results reveal that the anterior part of the tail in appendicularians could act as an anterior tail-mesoderm FGF source (Anterior Tail FGF Source, ATFS) of secreted Fgf9/16/20 ligands (**Figure 5B**). This may drastically differ with ascidians in which the Fgf9/16/20 source is restricted to the most posterior part of the tail (Posterior Tail FGF Source, PTFS), often associated with tail elongation and posterior cell identity differentiation or survival, similar to vertebrates (Diez del Corral and Storey 2004; Imai et al. 2004; Olivera-Martinez et al. 2012; Pasini et al. 2012). Interestingly, considering that the anterior region of the tail in ascidians serves as a source of RA signaling (Anterior Tail Retinoic Acid Source, ATRAS, **Figure 5B**) from the most anterior *Aldh1a*-positive cells of the tail muscle (Nagatomo and Fujiwara 2003), and given the antagonistic action between FGF and RA signaling conserved in ascidians and vertebrates, where down-regulation of RA signaling often leads to increased FGF production (Diez del Corral and Storey 2004; Olivera-Martinez et al. 2012; Pasini et al. 2012; Paschaki et al. 2013), it is tempting to speculate that the evolutionary innovation of this ATFS in appendicularians might be related to their loss of RA signaling (Cañestro and Postlethwait 2007; Martí-Solans et al. 2016) (**Figure 5**). This loss would have reduced selective constrains, allowing some *Fgf9/16/20* genes to gain novel expression domains in the anterior part of the tail. This drastic difference could represent a major shift in developmental signaling sources between appendicularians and ascidians, potentially driving the divergent evolution of developmental processes associated with the distinct body plans and lifestyles that characterize these two groups of tunicates.

The last examples of novel *Fgf* expression domains in *O. dioica* are related to the patterning of the epidermis in late hatchling stages. In the tail, the lateral wings that will develop into the tail fin express three *Fgf9/16/20* and two *Fgf11/12/13/14* paralogs. In the trunk, the oikoblast, which is the epidermal organ responsible for building the house, has recruited the expression of nearly all *Fgf9/16/20* and *Fgf11/12/13/14* paralogs, most of them in a broad fashion, although some appear to be restricted or excluded from certain fields within this complex organ (**Figure 4A11-J11**). The fact that no similar *Fgf* expression domains have been observed in the tail or the trunk of ascidian larvae (Satou et al. 2002; Imai et al. 2004) suggests that the recruitment of FGF for epidermis patterning could be an innovation of the appendicularian lineage, linked to the evolution of the house building organ and the tail movements that characterize its fully free-living lifestyle.

#### (4) extinction of ancestral expression domains linked to gene losses

In ascidians, *Fgf7/10/22* (which previously had been also referred as *Fgf3*) is strongly expressed throughout the ventral row of cells of the neural tube at tadpole stages, and its signaling function has been described to be crucial for the convergent extension of the notochord that underlies just underneath the neural tube (Shi et al. 2009). In *O. dioica*, the loss of *Fgf7/10/22* predicts that the convergent extension of the notochord might have become independent of the FGF signaling from the ventral neural tube, a prediction that can be tested in future experiments interfering with FGF signaling. Moreover, in ascidians *Fgf7/10/22* knockout makes tail absorption to be arrested during metamorphosis, which has led to suggest that *Fgf7/10/22* might play an inductive cue for the metamorphosis in ascidians (Treen et al. 2014). In this context, it is tempting to speculate that the loss of *Fgf7/10/22* in appendicularians could be related with the loss of a drastic metamorphosis and the lack of absorption of the tail, as it has been suggested in the ascidian-like biphasic tunicate ancestor (**Figure 5A**). Additionally, ascidian metamorphosis involves mesenchymal tissue, which consists of mesodermal cells that remain in a pluripotent state during development until postmetamorphic differentiation into adult tissues and structures. The ascidian *Fgf8/17/18* ortholog plays a role in the early differentiation of mesenchymal cells, with its expression maintained throughout embryonic development (Imai et al., 2004; Satou, 2020). Therefore, the loss of *Fgf8/17/18* appendicularians could be related to the loss of mesenchymal tissue and the lack of a drastic metamorphic process **(Figure 5)**. Altogether, our work highlights the evolution of the *Fgf* family in appendicularians as a paradigmatic example of what could be referred as “less, but more”, where massive gene losses, but also extensive duplications, result both in an overall conservation of *Fgf* expression domains, in many cases due to function shuffling among paralogs, and in the innovation of new expression domains. Interestingly, many of these innovations can be related to the transition from an ancestral ascidian-like biphasic lifestyle to the fully free-living lifestyle that characterizes appendicularians. Future functional analyses will be crucial to investigate several key aspects: the role of *Fgf11/12/13/14* gene on the innovation of a new mouth opening and the modulation of neural excitability; the involvement of multiple *Fgf* genes in the patterning of the oikoblast and the innovation of the house; the role of FGF signaling in notochord convergence and in the development of the fin; the impact of the emergence of an ATFS and its potential connection to the fact that these organisms are evolutionary knockouts of RA signaling; and finally, how the loss of *Fgf7/10/22* and *Fgf8/17/18* might be related to the loss of tail absorption and the absence of metamorphosis, both of which occurred during the evolutionary innovation of a fully free-living lifestyle of appendicularians.

## Acknowledgements

We thank present and past team members on CC’s laboratory for assistance and fruitful discussions on FGF signaling, gene loss, and evolution, specially to Sebastian Artime Paoletti for running the Oikopleura facility in the University of Barcelona. We thank to Centres Científics i Tecnològics de la UB for sea water supply and sequencing services.

## Author contributions

Conceptualization: CC, GS; formal Analysis: GS; funding acquisition: CC; investigation: GS, PB, JBR, AF, AFR, MF, NTA, JNW; methodology: GS, PB, JBR, AF, AFR, MF, NTA, RA; project administration: CC; resources: JNW, MJM, CP, NL; software: NTA, JNW, MJM, CP; CP supervision: CC; validation: GS, CC; visualization: GS, CC; writing (original draft): GS, CC; Writing (review & editing): GS, CC.

## Supplementary files

**Figure S1.**
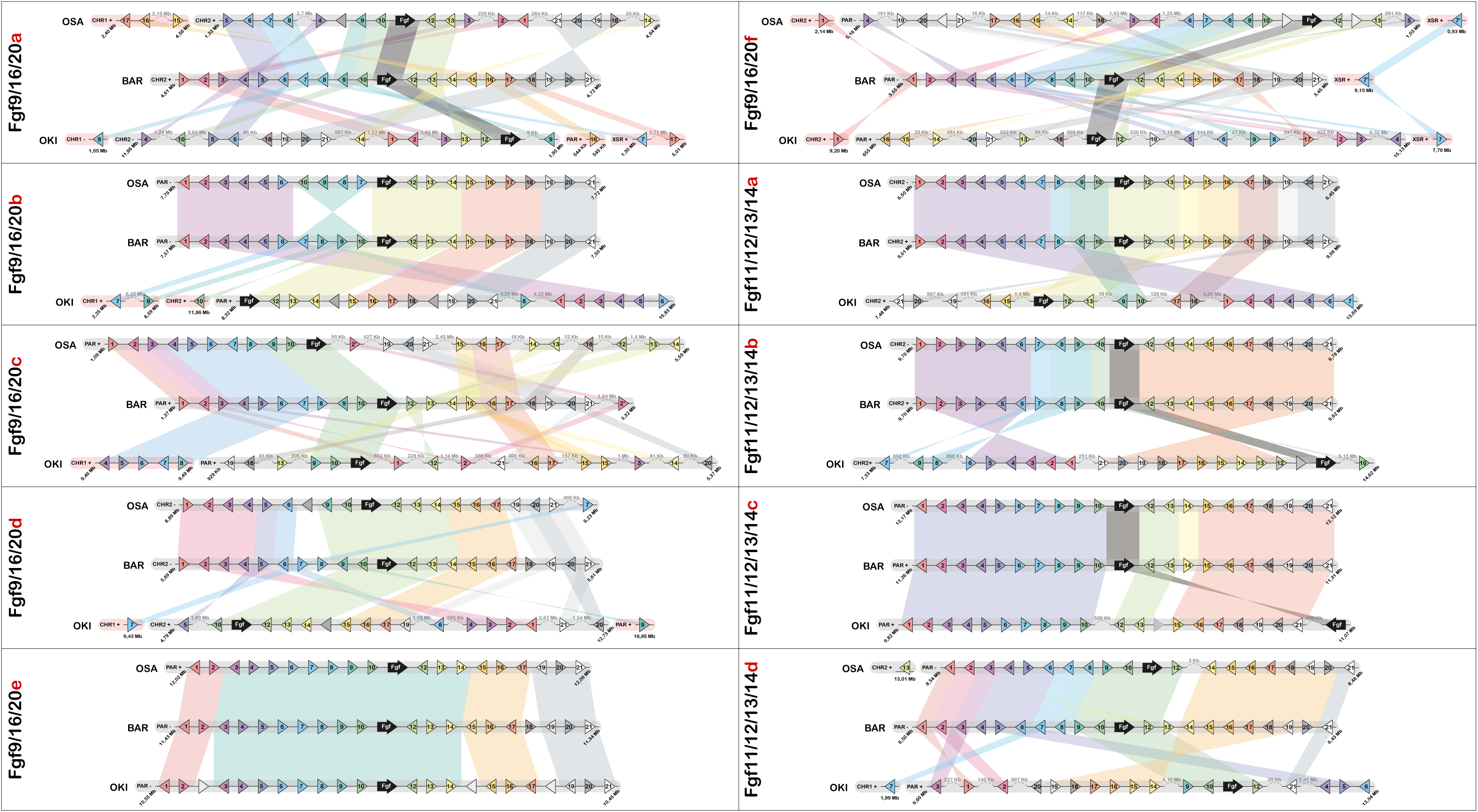
Comparative microsynteny conservation in the *Fgf* genes among the three *O. dioica* cryptic species. Comparison of microsynteny conservation between the genomic neighborhoods of Fgf genes (black arrow), and ten adjacent genes on each side. The BAR genome was used as the reference in comparisons with OSA and OKI.

**Figure S2.**
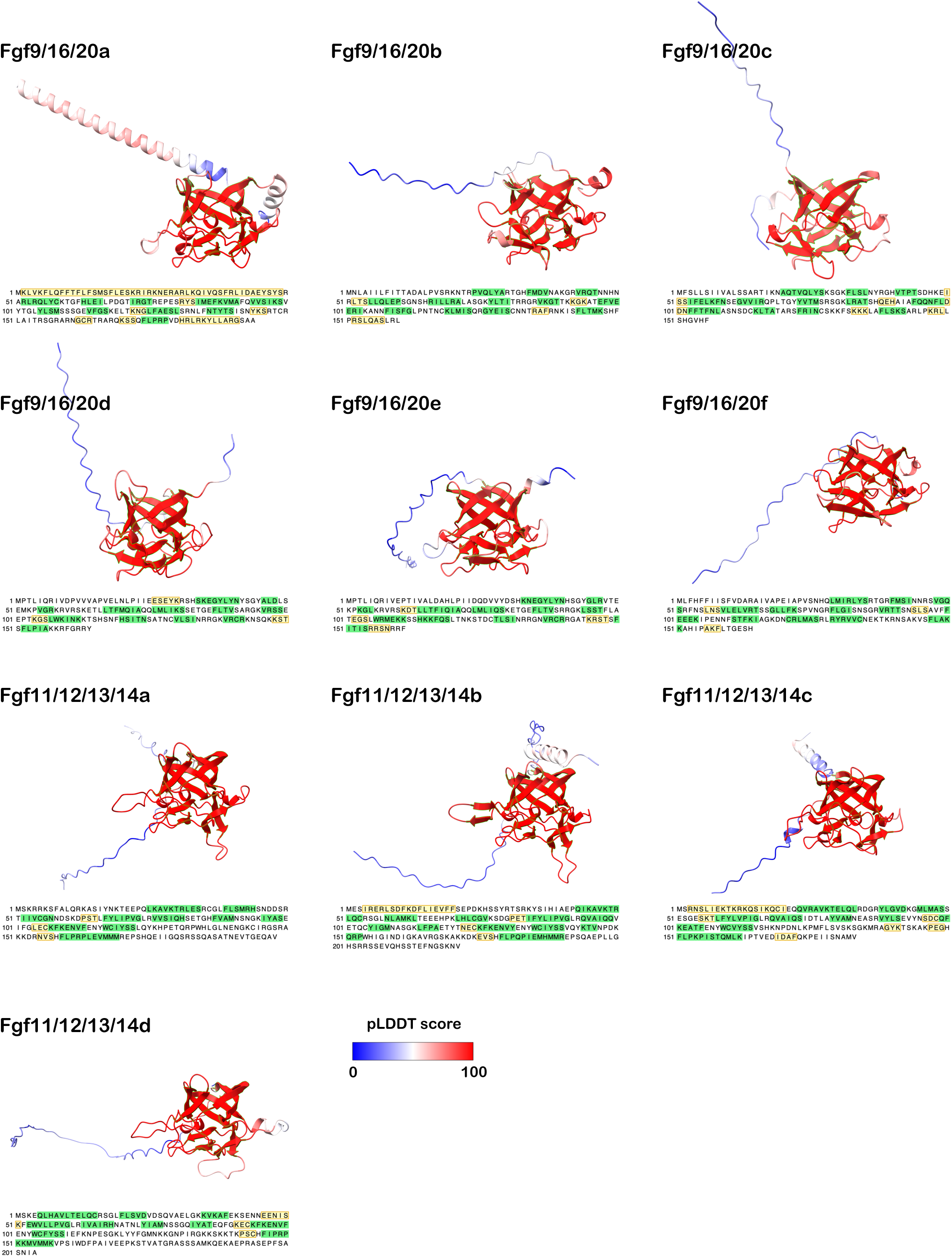
Three-dimensional models for *O. dioica* Fgf proteins. Models are colored according to their predicted local distance difference test (pLDDT) score. Predicted β-sheets are highlighted in light green in the models as well as in the protein sequences. Predicted α-helices are highlighted in light yellow in the protein sequences. All Fgf11/12/13/14 paralogs models and sequences correspond to the proximal isoform.

**Figure S3.**
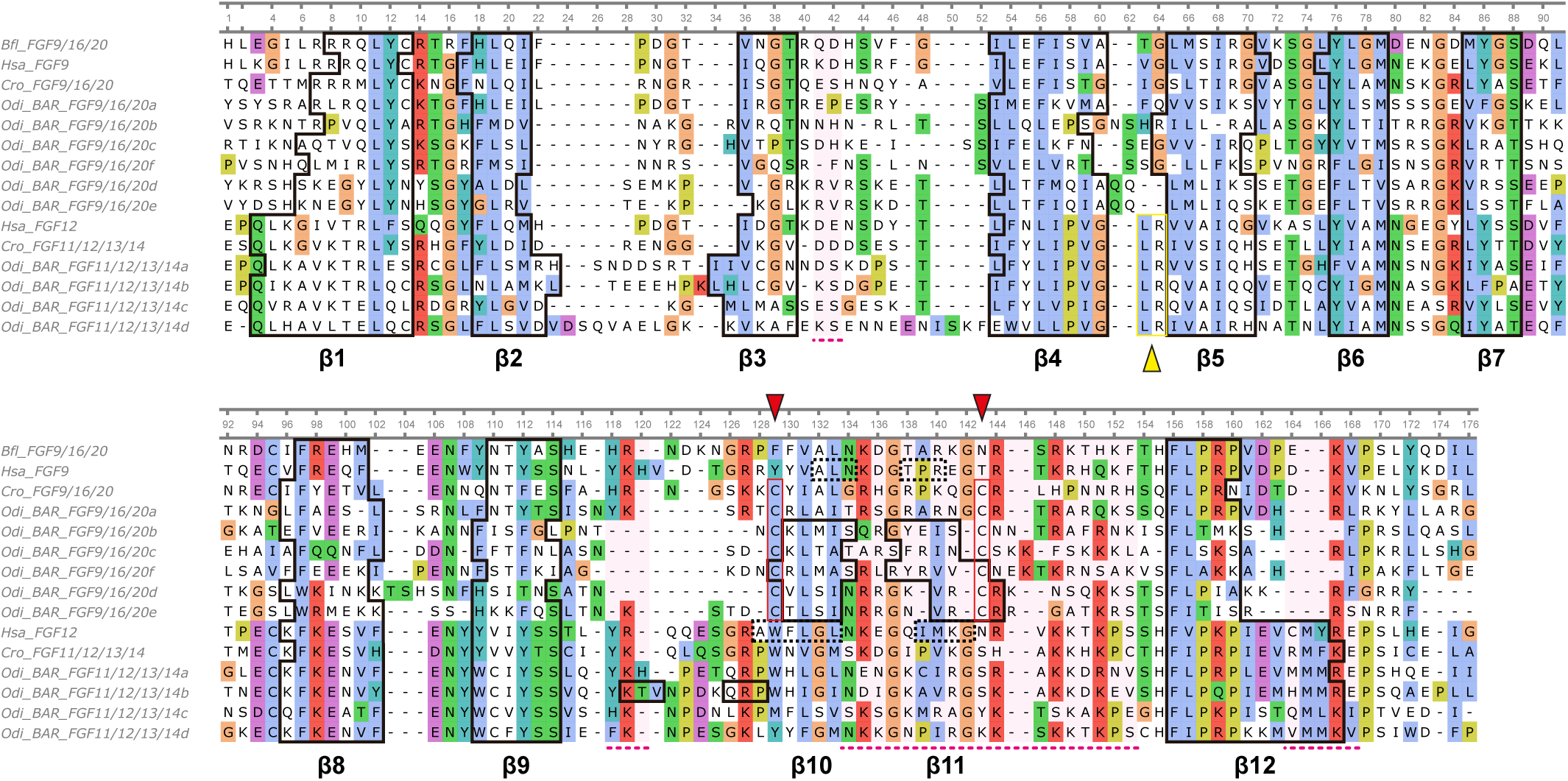
Protein alignment of *O. dioica* Fgf orthologs. The alignment includes the FGF domain of each ortholog. Sequence conservation is depicted according to the Clustal X default coloring. Black solid line boxes denote β-sheets as predicted by the AlphaFold2 software. Black dashed line boxes in Hsa_FGF9 and Hsa_FGF12 indicate β-sheets empirically confirmed but not predicted by AlphaFold2 (Goetz et al. 2009; Plotnikov et al. 2001). Red arrowheads and boxes in the *C. robusta* and *O. dioica* Fgf9/16/20 sequences highlight the positions of distinctive and conserved cysteines found in all tunicate Fgf9/16/20 paralogs. Yellow arrowheads and boxes in Fgf11/12/13/14 sequences denote the positions of the Leucine-Arginine pair, characteristic of the intracellular Fgf11/12/13/14 orthologs. Magenta shadings and dashed lines indicate the regions involved in binding heparin (Xu et al. 2012). Abbreviations: *Branchiostoma floridae* (Bfl), *Homo sapiens* (Hsa), *Ciona robusta* (Cro), *Oikopleura dioica* (Odi)

**Figure S4.**
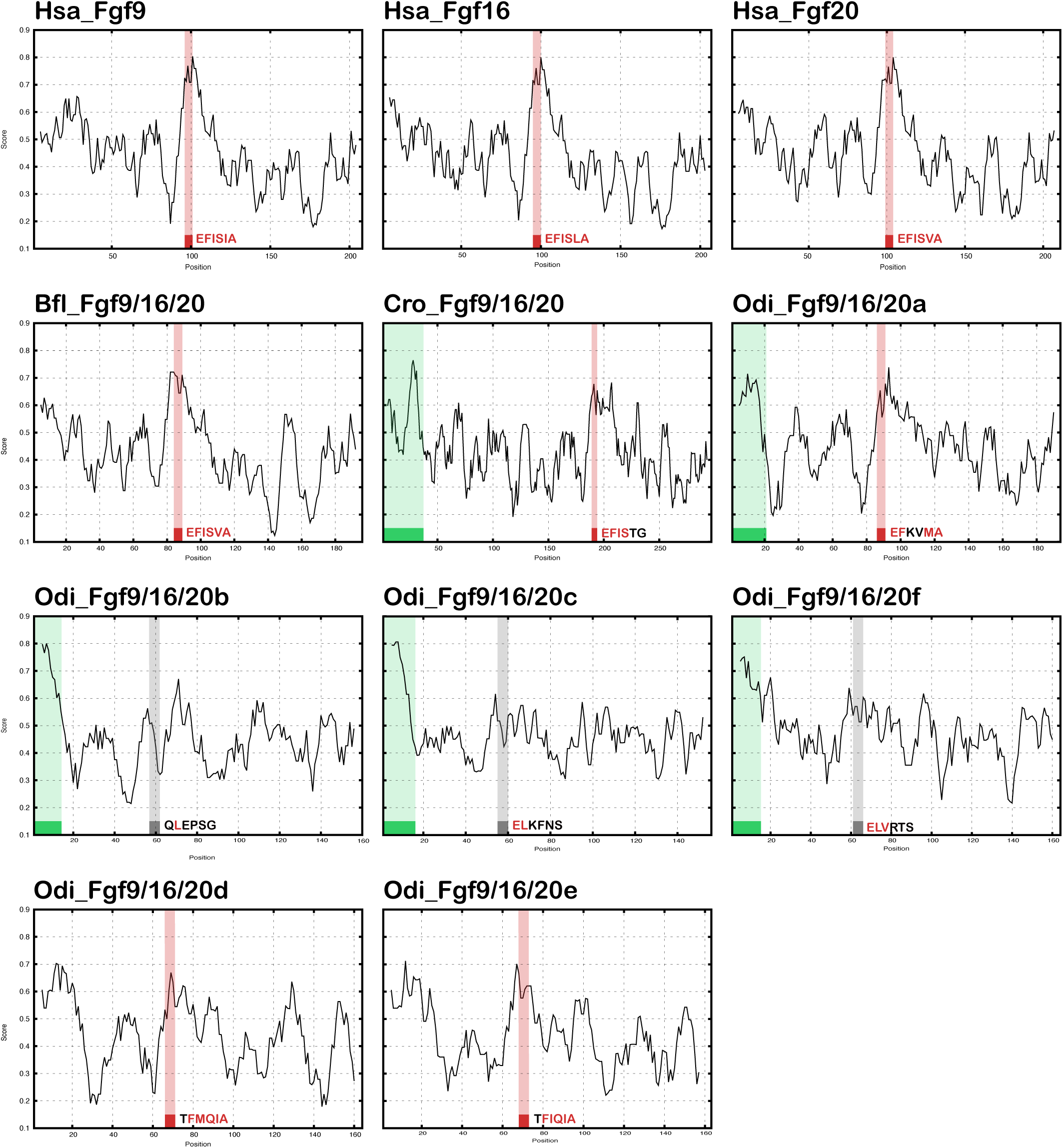
Hydropathy profile of chordate Fgf9/16/20 proteins. The graphs show the Kyte-Doolittle hydropathy score (y-axis) along the length of each protein sequence for *Homo sapiens* (Hsa), *Branchiostoma floridae* (Bfl), *Ciona robusta* (Cro), and *Oikopleura dioica* (Odi) Fgf9/16/20 orthologs. Scores are scaled to 1 for comparison purposes. Green squares and shaded areas indicate the presence and position of a cleavable signal peptide (SP) as predicted by Phobius. Red squares and shaded areas mark the position of well-conserved EFISIA motifs. Gray squares and shaded areas represent the positions of degraded EFISIA motifs. The amino acid letters in the EFISIA motif of each protein are colored red if they are conserved with the vertebrate counterpart or if they have been substituted by an amino acid with similar physicochemical properties.

**Table S1. Primers used for the cloning of *O. dioica Fgf* genes.**

**Table S2. Genomic loci of *Fgf* genes in the four sequenced genomes of *O. dioica*.**

**Table S3. List of sequences used for phylogenetic analyses.**

**Table S4. Fgf protein identity and similarity.**

**Supplementary file 1. Fgf protein alignment**

**Supplementary file 2. *Fgf* phylogenetic tree**

## Bibliography

1. Albalat R, Cañestro C. 2016. Evolution by gene loss. Nat Rev Genet 17:379–391.

2. Antoine M, Reimers K, Wirz W, Gressner AM, Müller R, Kiefer P. 2005. Fibroblast growth factor 3, a protein with a dual subcellular fate, is interacting with human ribosomal protein S2. Biochem Biophys Res Commun 338:1248–1255.

3. Bassham S, Cañestro C, Postlethwait JH. 2008. Evolution of developmental roles of Pax2/5/8 paralogs after independent duplication in urochordate and vertebrate lineages. BMC Biol 6:35.

4. Bassham S, Postlethwait J. 2000. Brachyury (T) expression in embryos of a larvacean urochordate, Oikopleura dioica, and the ancestral role of T. Dev Biol 220:322–32.

5. Bassham S, Postlethwait JH. 2005. The evolutionary history of placodes: a molecular genetic investigation of the larvacean urochordate Oikopleura dioica. Development 132:4259–4272.

6. Bertrand S, Aldea D, Oulion S, Subirana L, de Lera AR, Somorjai I, Escriva H. 2015. Evolution of the Role of RA and FGF Signals in the Control of Somitogenesis in Chordates. PLoS One 10:e0136587.

7. Bertrand S, Camasses A, Somorjai I, Belgacem MR, Chabrol O, Escande ML, Pontarotti P, Escriva H. 2011. Amphioxus FGF signaling predicts the acquisition of vertebrate morphological traits. Proc Natl Acad Sci U S A 108:9160–9165.

8. Bertrand S, Iwema T, Escriva H. 2014. FGF signaling emerged concomitantly with the origin of eumetazoans. Mol Biol Evol 31:310–318.

9. Bertrand V, Hudson C, Caillol D, Popovici C, Lemaire P. 2003. Neural tissue in ascidian embryos is induced by FGF9/16/20, acting via a combination of maternal GATA and Ets transcription factors. Cell 115:615–627.

10. Brozovic M, Dantec C, Dardaillon J, Dauga D, Faure E, Gineste M, Louis A, Naville M, Nitta KR, Piette J, et al. 2018. ANISEED 2017: Extending the integrated ascidian database to the exploration and evolutionary comparison of genome-scale datasets. Nucleic Acids Res 46:D718–D725.

11. Bryant DM, Stow JL. 2005. Nuclear Translocation of Cell-Surface Receptors: Lessons from Fibroblast Growth Factor. Traffic 6:947–953.

12. Cañestro C, Catchen JM, Rodríguez-Marí A, Yokoi H, Postlethwait JH. 2009. Consequences of lineage-specific gene loss on functional evolution of surviving paralogs: ALDH1A and retinoic acid signaling in vertebrate genomes. PLoS Genet 5:e1000496.

13. Cañestro C, Postlethwait JH. 2007. Development of a chordate anterior-posterior axis without classical retinoic acid signaling. Dev Biol 305:522–538.

14. Cañestro C, Yokoi H, Postlethwait JH. 2007. Evolutionary developmental biology and genomics. Nat Rev Genet 8:932–942.

15. Clarke T, Bouquet J-M, Fu X, Kallesøe T, Schmid M, Thompson EM. 2007. Rapidly evolving lamins in a chordate, Oikopleura dioica, with unusual nuclear architecture. Gene 396:159–169.

16. Coulier F, Pontarotti P, Roubin R, Hartung H, Goldfarb M, Birnbaum D. 1997. Of worms and men: an evolutionary perspective on the fibroblast growth factor (FGF) and FGF receptor families. J Mol Evol 44:43–56.

17. Crump JG, Maves L, Lawson ND, Weinstein BM, Kimmel CB. 2004. An essential role for Fgfs in endodermal pouch formation influences later craniofacial skeletal patterning. Development 131:5703–5716.

18. Czarkwiani A, Dylus D V., Carballo L, Oliveri P. 2021. FGF signalling plays similar roles in development and regeneration of the skeleton in the brittle star Amphiura filiformis. Development 148.

19. Danks G, Campsteijn C, Parida M, Butcher S, Doddapaneni H, Fu B, Petrin R, Metpally R, Lenhard B, Wincker P, et al. 2013. OikoBase: A genomics and developmental transcriptomics resource for the urochordate Oikopleura dioica. Nucleic Acids Res 41:1–9.

20. Dehal P, Boore JL. 2005. Two rounds of whole genome duplication in the ancestral vertebrate. PLoS Biol 3:e314.

21. Diez del Corral R, Storey KG. 2004. Opposing FGF and retinoid pathways: a signalling switch that controls differentiation and patterning onset in the extending vertebrate body axis. Bioessays 26:857–869.

22. Dorey K, Amaya E. 2010. FGF signalling: diverse roles during early vertebrate embryogenesis. Development 137:3731–3742.

23. Fan T-P, Su Y-H. 2015. FGF signaling repertoire of the indirect developing hemichordate Ptychodera flava. Mar Genomics 24:167–175.

24. Fan T-P, Ting H-C, Yu J-K, Su Y-H. 2018. Reiterative use of FGF signaling in mesoderm development during embryogenesis and metamorphosis in the hemichordate Ptychodera flava. BMC Evol Biol 18:120.

25. Fernández R, Gabaldón T. 2020. Gene gain and loss across the metazoan tree of life. Nat Ecol Evol 4:524–533.

26. Ferrández-Roldán A, Fabregà-Torrus M, Sánchez-Serna G, Duran-Bello E, Joaquín-Lluís M, Bujosa P, Plana-Carmona M, Garcia-Fernàndez J, Albalat R, Cañestro C. 2021. Cardiopharyngeal deconstruction and ancestral tunicate sessility. Nature 599:431–435.

27. Ferrández-Roldán A, Martí-Solans J, Cañestro C, Albalat R. 2019. Oikopleura dioica: An Emergent Chordate Model to Study the Impact of Gene Loss on the Evolution of the Mechanisms of Development. In: Tworzydlo W, Bilinski S, editors. Evo-Devo: Non-model Species in Cell and Developmental Biology. Vol. 68. Springer, Cham. p. 63–105.

28. Gasteiger E, Hoogland C, Gattiker A, Duvaud S, Wilkins MR, Appel RD, Bairoch A. 2005. Protein Identification and Analysis Tools on the ExPASy Server. In: The Proteomics Protocols Handbook. Totowa, NJ: Humana Press. p. 571–607.

29. Goldfarb M. 2001. Signaling By Fibroblast Growth Factors: The Inside Story. Science’s STKE 2001.

30. Goldfarb M. 2005. Fibroblast growth factor homologous factors: Evolution, structure, and function. Cytokine Growth Factor Rev 16:215–220.

31. Guijarro-Clarke C, Holland PWH, Paps J. 2020. Widespread patterns of gene loss in the evolution of the animal kingdom. Nat Ecol Evol 4:519–523.

32. Guindon S, Dufayard J-F, Lefort V, Anisimova M, Hordijk W, Gascuel O. 2010. New algorithms and methods to estimate maximum-likelihood phylogenies: assessing the performance of PhyML 3.0. Syst Biol 59:307–321.

33. Helsen J, Voordeckers K, Vanderwaeren L, Santermans T, Tsontaki M, Verstrepen KJ, Jelier R. 2020. Gene loss predictably drives evolutionary adaptation. Mol Biol Evol 37:2989–3002.

34. Hodgson JA, Pickrell JK, Pearson LN, Quillen EE, Prista A, Rocha J, Soodyall H, Shriver MD, Perry GH. 2014. Natural selection for the Duffy-null allele in the recently admixed people of Madagascar. Proc Biol Sci 281:20140930.

35. Horie R, Hazbun A, Chen K, Cao C, Levine M, Horie T. 2018. Shared evolutionary origin of vertebrate neural crest and cranial placodes. Nature 560:228–232.

36. Hudson C, Sirour C, Yasuo H. 2016. Co-expression of Foxa.a, Foxd and Fgf9/16/20 defines a transient mesendoderm regulatory state in ascidian embryos. Elife 5.

37. Imai KS, Hino K, Yagi K, Satoh N, Satou Y. 2004. Gene expression profiles of transcription factors and signaling molecules in the ascidian embryo: towards a comprehensive understanding of gene networks. Development 131:4047–4058.

38. Imai Kaoru S., Satoh N, Satou Y. 2002. Early embryonic expression of FGF4/6/9 gene and its role in the induction of mesenchyme and notochord in Ciona savignyi embryos. Development 129:1729– 1738.

39. Imai Karou S., Satoh N, Satou Y. 2002. Region specific gene expressions in the central nervous system of the ascidian embryo. Mech Dev 119 Suppl 1:S275–S277.

40. Imai KS, Stolfi A, Levine M, Satou Y. 2009. Gene regulatory networks underlying the compartmentalization of the Ciona central nervous system. Development 136:285–293.

41. Itoh N, Ornitz DM. 2011. Fibroblast growth factors: from molecular evolution to roles in development, metabolism and disease. J Biochem 149:121–130.

42. Jones P, Binns D, Chang H-Y, Fraser M, Li W, McAnulla C, McWilliam H, Maslen J, Mitchell A, Nuka G, et al. 2014. InterProScan 5: genome-scale protein function classification. Bioinformatics 30:1236–1240.

43. Jumper J, Evans R, Pritzel A, Green T, Figurnov M, Ronneberger O, Tunyasuvunakool K, Bates R, Žídek A, Potapenko A, et al. 2021. Highly accurate protein structure prediction with AlphaFold. Nature 596:583–589.

44. Käll L, Krogh A, Sonnhammer ELL. 2004. A combined transmembrane topology and signal peptide prediction method. J Mol Biol 338:1027–1036.

45. Kalyaanamoorthy S, Minh BQ, Wong TKF, von Haeseler A, Jermiin LS. 2017. ModelFinder: fast model selection for accurate phylogenetic estimates. Nat Methods 14:587–589.

46. Kim K, Gibboney S, Razy-Krajka F, Lowe EK, Wang W, Stolfi A. 2020. Regulation of Neurogenesis by FGF Signaling and Neurogenin in the Invertebrate Chordate Ciona. Front Cell Dev Biol 8:477.

47. Kirov A, Al-Hashimi H, Solomon P, Mazur C, Thorpe PE, Sims PJ, Tarantini F, Kumar TKS, Prudovsky I. 2012. Phosphatidylserine externalization and membrane blebbing are involved in the nonclassical export of FGF1. J Cell Biochem 113:956–966.

48. Kourakis MJ, Smith WC. 2007. A conserved role for FGF signaling in chordate otic/atrial placode formation. Dev Biol 312:245–257.

49. Krylov DM, Wolf YI, Rogozin IB, Koonin E V. 2003. Gene loss, protein sequence divergence, gene dispensability, expression level, and interactivity are correlated in eukaryotic evolution. Genome Res 13:2229–2235.

50. Laezza F, Lampert A, Kozel MA, Gerber BR, Rush AM, Nerbonne JM, Waxman SG, Dib-Hajj SD, Ornitz DM. 2009. FGF14 N-terminal splice variants differentially modulate Nav1.2 and Nav1.6-encoded sodium channels. Mol Cell Neurosci 42:90–101.

51. Lapraz F, Rottinger E, Duboc V, Range R, Duloquin L, Walton K, Wu SY, Bradham C, Loza MA, Hibino T, et al. 2006. RTK and TGF-beta signaling pathways genes in the sea urchin genome. Dev Biol 300:132–152.

52. Larsson A. 2014. AliView: A fast and lightweight alignment viewer and editor for large datasets. Bioinformatics 30:3276–3278.

53. Lu J, Wu T, Zhang B, Liu S, Song W, Qiao J, Ruan H. 2021. Types of nuclear localization signals and mechanisms of protein import into the nucleus. Cell Communication and Signaling 19:60.

54. Madeira F, Madhusoodanan N, Lee J, Eusebi A, Niewielska A, Tivey ARN, Lopez R, Butcher S. 2024. The EMBL-EBI Job Dispatcher sequence analysis tools framework in 2024. Nucleic Acids Res.

55. Martí-Solans J, Belyaeva O V., Torres-Aguila NP, Kedishvili NY, Albalat R, Cañestro C. 2016. Coelimination and Survival in Gene Network Evolution: Dismantling the RA-Signaling in a Chordate. Mol Biol Evol 33:2401–2416.

56. Martí-Solans J, Ferrández-Roldán A, Godoy-Marín H, Badia-Ramentol J, Torres-Aguila NP, Rodríguez-Marí A, Bouquet JM, Chourrout D, Thompson EM, Albalat R, et al. 2015. Oikopleura dioica culturing made easy: a low-cost facility for an emerging animal model in EvoDevo. Genesis 53:183–193.

57. Martí-Solans J, Godoy-Marín H, Diaz-Gracia M, Onuma TA, Nishida H, Albalat R, Cañestro C. 2021. Massive Gene Loss and Function Shuffling in Appendicularians Stretch the Boundaries of Chordate Wnt Family Evolution. Front Cell Dev Biol 9:700827.

58. Matus DQ, Thomsen GH, Martindale MQ. 2007. FGF signaling in gastrulation and neural development in Nematostella vectensis, an anthozoan cnidarian. Dev Genes Evol 217:137–148.

59. McClintock JM, Carlson R, Mann DM, Prince VE. 2001. Consequences of Hox gene duplication in the vertebrates: an investigation of the zebrafish Hox paralogue group 1 genes. Development 128:2471–2484.

60. Mikhaleva Y, Skinnes R, Sumic S, Thompson EM, Chourrout D. 2018. Development of the house secreting epithelium, a major innovation of tunicate larvaceans, involves multiple homeodomain transcription factors. Dev Biol 443:117–126.

61. Miyakawa K, Hatsuzawa K, Kurokawa T, Asada M, Kuroiwa T, Imamura T. 1999. A hydrophobic region locating at the center of fibroblast growth factor-9 is crucial for its secretion. J Biol Chem 274:29352–29357.

62. Miyakawa K, Imamura T. 2003. Secretion of FGF-16 requires an uncleaved bipartite signal sequence. Journal of Biological Chemistry 278:35718–35724.

63. Miyazaki Y, Nishida H, Kumano G. 2007. Brain induction in ascidian embryos is dependent on juxtaposition of FGF9 / 16 / 20-producing and -receiving cells.

64. Munoz-Sanjuan I, Smallwood PM, Nathans J. 2000. Isoform diversity among fibroblast growth factor homologous factors is generated by alternative promoter usage and differential splicing. J Biol Chem 275:2589–2597.

65. Nagatomo K, Fujiwara S. 2003. Expression of Raldh2, Cyp26 and Hox-1 in normal and retinoic acid-treated Ciona intestinalis embryos. Gene Expr Patterns 3:273–277.

66. Naville M, Henriet S, Warren I, Sumic S, Reeve M, Volff JN, Chourrout D. 2019. Massive Changes of Genome Size Driven by Expansions of Non-autonomous Transposable Elements. Current Biology 29:1161–1168.

67. Nguyen Ba AN, Pogoutse A, Provart N, Moses AM. 2009. NLStradamus: a simple Hidden Markov Model for nuclear localization signal prediction. BMC Bioinformatics 10:202.

68. Nguyen L-T, Schmidt HA, von Haeseler A, Minh BQ. 2015. IQ-TREE: A Fast and Effective Stochastic Algorithm for Estimating Maximum-Likelihood Phylogenies. Mol Biol Evol 32:268–274.

69. Nishida H. 2008. Development of the appendicularian Oikopleura dioica: culture, genome, and cell lineages. Dev Growth Differ 50 Suppl 1:S239–S256.

70. Novembre J, Galvani AP, Slatkin M. 2005. The geographic spread of the CCR5 Delta32 HIV-resistance allele. PLoS Biol 3:e339.

71. Olivera-Martinez I, Harada H, Halley PA, Storey KG. 2012. Loss of FGF-Dependent Mesoderm Identity and Rise of Endogenous Retinoid Signalling Determine Cessation of Body Axis Elongation. PLoS Biol 10:e1001415.

72. Olsen LC, Kourtesis I, Busengdal H, Jensen MF, Hausen H, Chourrout D. 2018. Evidence for a centrosome-attracting body like structure in germ-soma segregation during early development, in the urochordate Oikopleura dioica. BMC Dev Biol 18:4.

73. Olsen SK, Garbi M, Zampieri N, Eliseenkova A V., Ornitz DM, Goldfarb M, Mohammadi M. 2003. Fibroblast Growth Factor (FGF) Homologous Factors Share Structural but Not Functional Homology with FGFs. Journal of Biological Chemistry 278:34226–34236.

74. Olsnes S, Klingenberg O, Więdłocha A. 2003. Transport of Exogenous Growth Factors and Cytokines to the Cytosol and to the Nucleus. Physiol Rev 83:163–182.

75. Olson M V. 1999. When less is more: gene loss as an engine of evolutionary change. Am J Hum Genet 64:18–23.

76. Ornitz DM, Itoh N. 2015. The fibroblast growth factor signaling pathway. Wiley Interdiscip Rev Dev Biol 4:215–266.

77. Ornitz DM, Itoh N. 2022. New developments in the biology of fibroblast growth factors. WIREs mechanisms of disease 14:e1549.

78. Osipova E, Barsacchi R, Brown T, Sadanandan K, Gaede AH, Monte A, Jarrells J, Moebius C, Pippel M, Altshuler DL, et al. 2023. Loss of a gluconeogenic muscle enzyme contributed to adaptive metabolic traits in hummingbirds. Science 379:185–190.

79. Oulion S, Bertrand S, Escriva H. 2012. Evolution of the FGF Gene Family. Int J Evol Biol 2012:1–12.

80. Owji H, Nezafat N, Negahdaripour M, Hajiebrahimi A, Ghasemi Y. 2018. A comprehensive review of signal peptides: Structure, roles, and applications. Eur J Cell Biol 97:422–441.

81. Pablo JL, Pitt GS. 2016. Fibroblast Growth Factor Homologous Factors: New Roles in Neuronal Health and Disease. Neuroscientist 22:19–25.

82. Paschaki M, Schneider C, Rhinn M, Thibault-Carpentier C, Dembélé D, Niederreither K, Dollé P. 2013. Transcriptomic analysis of murine embryos lacking endogenous retinoic acid signaling. PLoS One 8:e62274.

83. Pasini A, Manenti R, Rothbacher U, Lemaire P. 2012. Antagonizing retinoic acid and FGF/MAPK pathways control posterior body patterning in the invertebrate chordate Ciona intestinalis. PLoS One 7:e46193.

84. Pettersen EF, Goddard TD, Huang CC, Meng EC, Couch GS, Croll TI, Morris JH, Ferrin TE. 2021. UCSF ChimeraX: Structure visualization for researchers, educators, and developers. Protein Sci 30:70–82.

85. Plessy C, Mansfield MJ, Bliznina A, Masunaga A, West C, Tan Y, Liu AW, Grašič J, del Río Pisula MS, Sánchez-Serna G, et al. 2024. Extreme genome scrambling in marine planktonic *Oikopleura dioica* cryptic species. Genome Res.

86. Plotnikov AN, Eliseenkova A V., Ibrahimi OA, Shriver Z, Sasisekharan R, Lemmon MA, Mohammadi M. 2001. Crystal Structure of Fibroblast Growth Factor 9 Reveals Regions Implicated in Dimerization and Autoinhibition. Journal of Biological Chemistry 276:4322–4329.

87. Popovici C, Conchonaud F, Birnbaum D, Roubin R. 2004. Functional Phylogeny Relates LET-756 to Fibroblast Growth Factor 9. Journal of Biological Chemistry 279:40146–40152.

88. Popovici C, Fallet M, Marguet D, Birnbaum D, Roubin R. 2006. Intracellular trafficking of LET-756, a fibroblast growth factor of C. elegans, is controlled by a balance of export and nuclear signals. Exp Cell Res 312:1484–1495.

89. Popovici C, Roubin R, Coulier F, Birnbaum D. 2005. An evolutionary history of the FGF superfamily. Bioessays 27:849–857.

90. Rees JM, Palmer MA, Gillis JA. 2024. Fgf signalling is required for gill slit formation in the skate, Leucoraja erinacea. Dev Biol 506:85–94.

91. Revest JM, DeMoerlooze L, Dickson C. 2000. Fibroblast growth factor 9 secretion is mediated by a non-cleaved amino-terminal signal sequence. Journal of Biological Chemistry 275:8083–8090.

92. Röttinger E, Saudemont A, Duboc V, Besnardeau L, McClay D, Lepage T. 2008. FGF signals guide migration of mesenchymal cells, control skeletal morphogenesis [corrected] and regulate gastrulation during sea urchin development. Development 135:353–365.

93. Satou Y. 2020. A gene regulatory network for cell fate specification in Ciona embryos. Curr Top Dev Biol 139:1–33.

94. Satou Y, Imai KS, Satoh N. 2002. Fgf genes in the basal chordate Ciona intestinalis. Dev Genes Evol 212:432–8.

95. Schäfer T, Zentgraf H, Zehe C, Brügger B, Bernhagen J, Nickel W. 2004. Unconventional Secretion of Fibroblast Growth Factor 2 Is Mediated by Direct Translocation across the Plasma Membrane of Mammalian Cells. Journal of Biological Chemistry 279:6244–6251.

96. Schlessinger J, Plotnikov AN, Ibrahimi OA, Eliseenkova A V., Yeh BK, Yayon A, Linhardt RJ, Mohammadi M. 2000. Crystal Structure of a Ternary FGF-FGFR-Heparin Complex Reveals a Dual Role for Heparin in FGFR Binding and Dimerization. Mol Cell 6:743–750.

97. Schoorlemmer J, Goldfarb M. 2002. Fibroblast Growth Factor Homologous Factors and the Islet Brain-2 Scaffold Protein Regulate Activation of a Stress-activated Protein Kinase. Journal of Biological Chemistry 277:49111–49119.

98. Sharma V, Hecker N, Roscito JG, Foerster L, Langer BE, Hiller M. 2018. A genomics approach reveals insights into the importance of gene losses for mammalian adaptations. Nat Commun 9:1215.

99. Sheng Z, Liang Y, Lin C-Y, Comai L, Chirico WJ. 2005. Direct Regulation of rRNA Transcription by Fibroblast Growth Factor 2. Mol Cell Biol 25:9419–9426.

100. Shi W, Peyrot SM, Munro E, Levine M. 2009. FGF3 in the floor plate directs notochord convergent extension in the Ciona tadpole. Development 136:23–28.

101. Sluzalska KD, Slawski J, Sochacka M, Lampart A, Otlewski J, Zakrzewska M. 2021. Intracellular partners of fibroblast growth factors 1 and 2 - implications for functions. Cytokine Growth Factor Rev 57:93–111.

102. Smallwood PM, Munoz-Sanjuan I, Tong P, Macke JP, Hendry SH, Gilbert DJ, Copeland NG, Jenkins NA, Nathans J. 1996. Fibroblast growth factor (FGF) homologous factors: new members of the FGF family implicated in nervous system development. Proc Natl Acad Sci U S A 93:9850–9857.

103. Stach T, Winter J, Bouquet J-MM, Chourrout D, Schnabel R. 2008. Embryology of a planktonic tunicate reveals traces of sessility. Proc Natl Acad Sci U S A 105:7229–7234.

104. Stolfi A, Ryan K, Meinertzhagen IA, Christiaen L. 2015. Migratory neuronal progenitors arise from the neural plate borders in tunicates. Nature 527:371–374.

105. Technau U, Scholz CB. 2003. Origin and evolution of endoderm and mesoderm. Int J Dev Biol 47:531– 539.

106. Teufel F, Almagro Armenteros JJ, Johansen AR, Gíslason MH, Pihl SI, Tsirigos KD, Winther O, Brunak S, von Heijne G, Nielsen H. 2022. SignalP 6.0 predicts all five types of signal peptides using protein language models. Nat Biotechnol 40:1023–1025.

107. Teven CM, Farina EM, Rivas J, Reid RR. 2014. Fibroblast growth factor (FGF) signaling in development and skeletal diseases. Genes Dis 1:199–213.

108. Thisse B, Thisse C. 2005. Functions and regulations of fibroblast growth factor signaling during embryonic development. Dev Biol 287:390–402.

109. Treen N, Chavarria E, Weaver CJ, Brangwynne CP, Levine M. 2023. An FGF timer for zygotic genome activation. Genes Dev 37:80–85.

110. Treen N, Yoshida K, Sakuma T, Sasaki H, Kawai N, Yamamoto T. 2014. Tissue-specific and ubiquitous gene knockouts by TALEN electroporation provide new approaches to investigating gene function in Ciona.: 481–487.

111. Veeman MT, Newman-Smith E, El-Nachef D, Smith WC. 2010. The ascidian mouth opening is derived from the anterior neuropore: reassessing the mouth/neural tube relationship in chordate evolution. Dev Biol 344:138–149.

112. Wagner E, Levine M. 2012. FGF signaling establishes the anterior border of the Ciona neural tube. Development 139:2351–2359.

113. Wang Chaojian, Wang Chuan, Hoch EG, Pitt GS. 2011. Identification of novel interaction sites that determine specificity between fibroblast growth factor homologous factors and voltage-gated sodium channels. J Biol Chem 286:24253–24263.

114. Wang K, Omotezako T, Kishi K, Nishida H, Onuma TA. 2015. Maternal and zygotic transcriptomes in the appendicularian, Oikopleura dioica: novel protein-encoding genes, intra-species sequence variations, and trans-spliced RNA leader. Dev Genes Evol 225:149–159.

115. Wilson V, Olivera-Martinez I, Storey KG. 2009. Stem cells, signals and vertebrate body axis extension. Development 136:1591–1604.

116. Wu Q-F, Yang L, Li S, Wang Q, Yuan X-B, Gao X, Bao L, Zhang X. 2012. Fibroblast Growth Factor 13 Is a Microtubule-Stabilizing Protein Regulating Neuronal Polarization and Migration. Cell 149:1549–1564.

117. Xie Y, Su N, Yang J, Tan Q, Huang S, Jin M, Ni Z, Zhang B, Zhang D, Luo F, et al. 2020. FGF/FGFR signaling in health and disease. Signal Transduct Target Ther 5:181.

118. Xu R, Ori A, Rudd TR, Uniewicz KA, Ahmed YA, Guimond SE, Skidmore MA, Siligardi G, Yates EA, Fernig DG. 2012. Diversification of the structural determinants of fibroblast growth factor-heparin interactions: implications for binding specificity. J Biol Chem 287:40061–40073.

119. Xu Y-C, Guo Y-L. 2020. Less Is More, Natural Loss-of-Function Mutation Is a Strategy for Adaptation. Plant Commun 1:100103.

120. Yasuo H, Hudson C. 2007. FGF8/17/18 functions together with FGF9/16/20 during formation of the notochord in Ciona embryos. Dev Biol 302:92–103.

